# Spatially resolved cellular and circuit architecture of the insular cortex controlling different dimensions of pain

**DOI:** 10.64898/2026.07.15.738697

**Authors:** Guoguang Xie, Meng Wang, Yiqiong Liu, Yi Zhang

## Abstract

Chronic pain is a leading cause of human suffering, affecting about 20% of the population. The insular cortex (IC) is a central hub for the multidimensional experience of pain, yet how the diverse neurons in IC are organized and assembled into functional circuits remains unclear. Here, combining multiplexed error-robust fluorescence *in situ* hybridization (MERFISH), neural tracing and *in vivo* functional analyses, we resolve the molecular, cellular and circuit architecture of the IC. We find that neuronal projections are specified by both transcriptomic identity and spatial topography. We further show that three molecularly defined neuron types in the posterior IC form discrete brain-wide networks that differentially regulate pain. Layer 5 pyramidal tract (PT) neurons regulate mechanical, thermal and affective pain. In contrast, layer 5 intratelencephalic (IT) neurons selectively modulate thermal nociception, whereas layer 6 corticothalamic (CT) neurons surprisingly relieve negative pain affects. Together, these findings reveal the spatially resolved cellular and circuit organization of the IC governing different dimensions of pain and uncover targets for precision pain therapy.

## Introduction

Neurons exhibit extraordinary molecular, anatomical, and functional diversity and are highly organized spatially (*1–4*). A prevailing hypothesis posits that the transcriptome of a neuron encodes its developmental trajectory, thereby dictating its morphology, connectivity, and function(*2, 5*). However, the fidelity of such transcriptomic-to-phenotypic correlation varies substantially. While robust correspondence exists in the retinal and olfactory neurons(*6–8*), such one-to-one mappings are rare in the mammalian cerebral cortex and hypothalamus(*9–11*). Instead, cortical neurons sharing a homogeneous transcriptomic profile often display highly variable morphologies, long-range projections and functional properties(*2, 12*). Recent advances in spatial transcriptomics enable the mapping of this architecture, revealing overlapping spatial distributions of some transcriptomically defined projecting neurons in the mouse motor cortex(*13*). Furthermore, although spatial position interacts with the transcriptome to define neuronal phenotype in the zebrafish optic tectum(*14*), it remains unknown whether similar topographical principles apply to mammalian cerebral cortex. Consequently, deciphering how transcriptomic identity, spatial organization, and projection patterns integrate to support cortical computation and behavioral functions remains a critical challenge.

The insular cortex (IC) comprises diverse cell types and is extensively connected across the brain(*15–17*), representing an ideal system to decode the correspondence between transcriptomic identity, spatial organization and connectivity. Importantly, the IC is well positioned as a critical hub for pain processing, orchestrating multimodal sensory information, emotion and cognition(*18–21*). Clinical evidence suggests that the IC is one of the most consistently activated regions under pain conditions, whereas maladaptive plasticity in the IC and disrupted connectivity with the limbic network are frequently observed in patients with chronic pain(*21–23*). Notably, direct electrical stimulation of the IC elicits pain sensations(*24*), demonstrating the critical roles of IC in pain perception. Preclinical studies show that the IC integrates nociceptive inputs and modulates both sensory and affective dimensions of pain through projections to the amygdala, thalamus, and descending pathways including the rostral ventromedial medulla (RVM) or directly to the spinal cord(*25–28*). Indeed, optogenetic activation of the IC drives hyperalgesia and aversion, whereas inhibition of the IC alleviates hypersensitivity and anxiety-like behaviors in mice(*26*). However, the IC is highly heterogeneous and the cellular basis in IC underlying the critical functions in pain processing remains poorly defined. While layer-specific projections in the motor cortex regulate distinct pain dimensions(*29*), the organizational principles of the IC in pain processing remain largely unknown. In particular, it is unclear how molecularly distinct cell types are spatially organized and wired into different functional circuits, thereby regulating different dimensions of pain. Moreover, how these cell types are affected in chronic pain and whether their manipulation can alleviate pain remain unresolved.

Here we used MERFISH to decode the spatial organization of the IC and adjacent regions, revealing distinct patterns of neuronal subtype distribution and gene expression. We then integrated MERFISH with retrograde tracing to determine how transcriptomic identity, spatial distribution and projection patterns covary across the tissue. Our data showed that, although major neuron classes exhibit highly distinct projection patterns, individual transcriptomic types within the same class share broadly overlapping projection targets with specific topographic preferences. We further showed that molecularly defined neuron types are differentially integrated into the neural networks, respond distinctly to noxious stimulation and play separable roles in pain processing. Together, our results provide a multimodal framework for understanding the spatially resolved molecular, cellular and circuit architecture of the IC and reveal discrete neural substrates that regulate different dimensions of pain.

## Results

### MERFISH profiling of the mouse insular cortex

To study the neural basis of the mouse insular cortex (IC) in pain processing, we performed MERFISH, a method for single-cell spatial transcriptomics, on coronal sections containing the IC and adjacent regions to decode the cellular heterogeneity and spatial organization (**Fig. 1A**). We selected a panel of 434 genes for MERFISH imaging, including cell-type markers, receptors, ion channels, and activity-related genes (ARGs) (**table S1-S3**; see Methods for details). We analyzed 16 serial coronal sections of a male mouse, spanning the anterior-posterior (AP) axis of the IC at approximately 200 µm intervals (Bregma +2.0 to -1.2 mm; **Fig. 1B**). Following RNA transcript detection and cell segmentation (Methods), we obtained 103,085 high quality cells (**fig. S1A**) (**table S4**). We performed MERFISH with another replicate containing 8 serial coronal sections at 400 µm intervals and obtained 43,281 high quality single cells (**fig. S1B**).

**Fig. 1.**
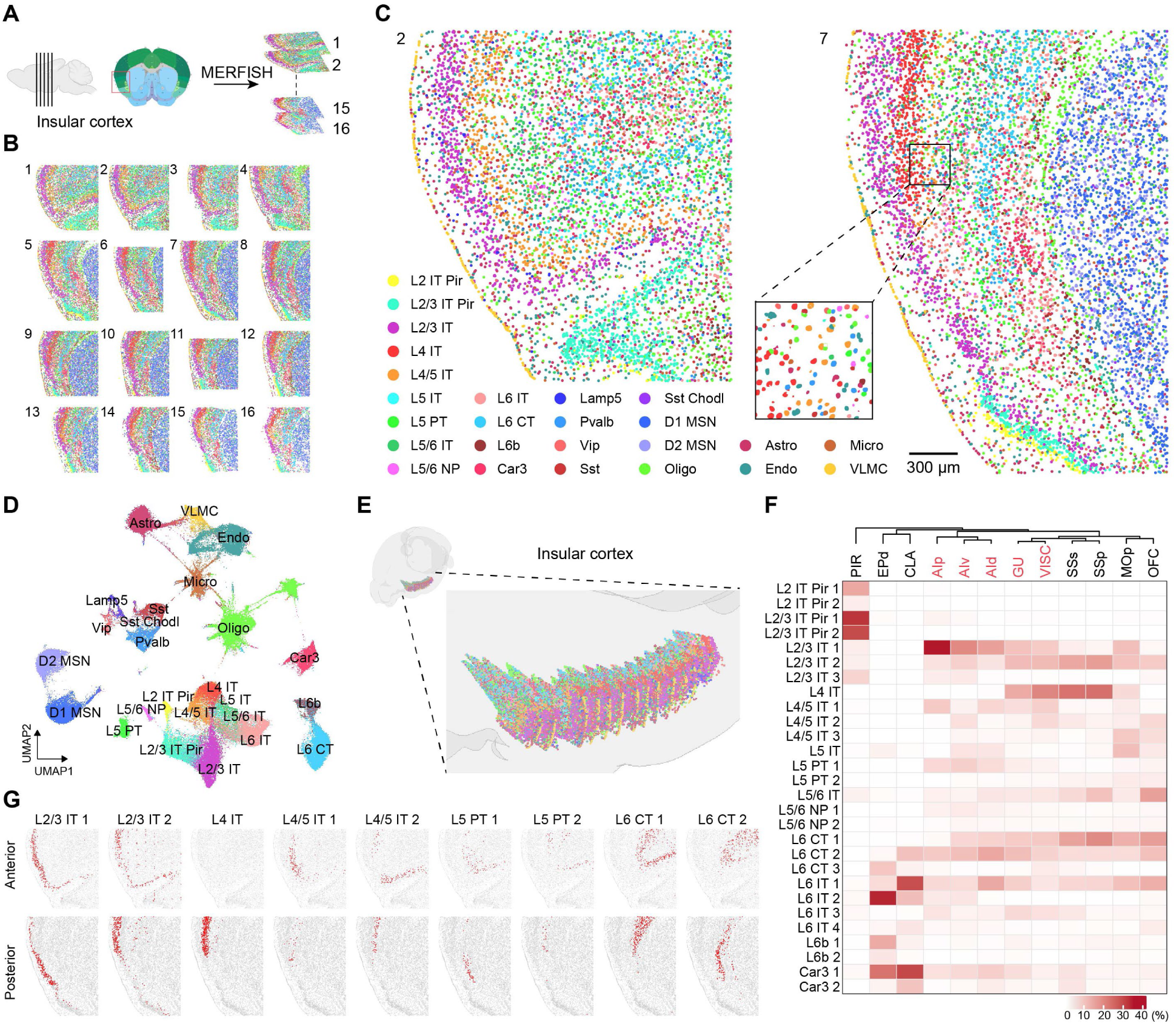
Spatial organization of the molecularly distinct cell types in the IC and adjacent regions. **(A)** Diagram of serial sections with the IC for MERFISH analysis. **(B)** An overview of all spatial single-cell maps of the cell types in the 16 coronal slices of a mouse spanning anterior to posterior IC. Cells are color-coded by their identities. **(C)** Spatial maps of the cell types in two representative coronal slices for anterior (left) and posterior (right) parts of the IC, respectively. An enlarged view of the segmented cells for the indicated region is shown in the insert. Scale bar, 300 μm. **(D)** UMAP visualization of all cells integrating two male mice replicates identified by MERFISH (total number of cells = 146,366). **(E)** 3D reconstruction of all the cells in the IC of a male mouse after registered to the Allen CCFv3. **(F)** Heatmap showing the proportion of excitatory neuron subtypes in different anatomical regions. The subregions of IC are highlighted in red. **(G)** Spatial location of nine representative excitatory neuron subtypes on the representative coronal slices for anterior (top) and posterior (bottom) parts of IC. Red dots represent the indicated subtypes.

We identified 25 transcriptomically distinct cell populations in the IC and adjacent regions by unsupervised clustering analysis and revealed their spatial organization (**Fig. 1, C and D**). These populations spanned three major families: excitatory neurons, inhibitory neurons, and non-neuronal cells. Excitatory neurons comprised canonical cortical classes, including intratelencephalic (IT), near-projecting (NP), pyramidal tract (PT), and cortico-thalamic (CT) neurons at specific layers (L). Inhibitory neurons included Vip, Lamp5, Pvalb, Sst, and Sst Chodl populations, whereas non-neuronal cells included oligodendrocytes (Oligo), astrocytes (Astro), microglia (Micro), endothelial cells (Endo), and vascular leptomeningeal cells (VLMC) (**Fig. 1D and fig. S1C**). The cellular classes identified by MERFISH were highly reproducible across biological replicates (**fig. S1D**) and showed strong correspondence with those identified by single-cell RNA sequencing (scRNA-seq) from public data(*30*) (**fig. S1E**). Fine-grained clustering further resolved these classes into 28 excitatory, 8 inhibitory, and 7 non-neuronal transcriptomic types (**fig. S2A-C**). Excitatory neuronal types exhibited a pronounced laminar organization, a characteristic of the cerebral cortex, with molecularly distinct populations segregating into specific layers, whereas molecularly similar neurons were localized within the same cortical layer (**Fig. 1, B and C and fig. S2B**). In contrast, inhibitory neurons and most non-neuronal cells were distributed broadly across all layers, though oligodendrocytes were enriched near fiber tracts and VLMCs localized along the pial surface (**Fig. 1, B and C and fig. S2B**). Finally, to assess the potential sex-specific differences in cellular architecture, we profiled MERFISH analysis on the IC sections of a female mouse (**fig. S2D**). Comparative analysis revealed no detectable differences in cellular composition between male and female mice, as indicated by UMAP analysis (**fig. S2, E and F**).

### Cellular composition and spatial organization of the mouse insular cortex

To investigate the spatial architecture of cell types in the IC and adjacent areas, we registered our MERFISH images to the Allen mouse brain Common Coordinate Framework version 3 (Allen CCFv3) using the decoded spatial maps (**fig. S3A**). This enabled us to reconstruct the 3D organization of the entire imaged regions (**fig. S3B**) and the IC subregions specifically (**Fig. 1E**). Notably, excitatory neurons in the claustrum (CLA) exhibited transcriptomic profiles highly similar to the excitatory neuronal types in the deep layers of the IC(*31*) (**fig. S3C**). Following anatomical annotation based on the Allen CCFv3 registration, we quantified the cellular composition across the IC subregions and revealed the excitatory neurons as the largest population, among which IT neurons are the most abundant types (**fig. S3D**). Along the AP axis, major cell classes and neuron types exhibited broad spatial distributions, although several populations displayed localized enrichment within restricted AP domains (**fig. S3, E and F**). For example, L4 IT neurons were absent from anterior sections, L4/5 IT neurons were less abundant in medial sections, and Car3 neurons were medially enriched, whereas L5 PT neurons were uniformly distributed along the AP axis (**fig. S3E**). Fine-grained analysis revealed distinct AP spatial biases among specific neuron types. For example, the L2/3 IT 2, L5 IT, L6 IT 1, and L6 CT 1 neurons were enriched in anterior sections, whereas L2/3 IT 1, L4 IT, L6 IT 2, and L6 IT 3 were more abundant in posterior sections (**fig. S3F**). Together, these results revealed the cellular composition of the IC and the spatial organization of the cell types along the AP axis.

To further define the spatial organization of the IC and adjacent areas, we performed spatial niche analysis based on gene expression profiles and found prominent laminar niches in both anterior and posterior sections (**fig. S4A**), further supporting the layered organization. Based on anatomical annotation with Allen CCFv3, we next quantified the subregional distribution of excitatory (**Fig. 1F**) and inhibitory neuron types (**fig. S4B**). Given the transcriptomic similarity between CLA neurons and IC neurons (**fig. S3C**), we applied cortical nomenclature to both structures for simplicity. The paleocortical and subcortical regions, including the piriform cortex (Pir), dorsal endopiriform nucleus (EPd), and CLA, exhibited distinct cellular compositions compared to neocortical regions (**Fig. 1F**). Specifically, Pir was dominated by Pir-specific excitatory neurons, whereas EPd and CLA contained subtypes that are molecularly congruent with neurons in IC deep layers (**Fig. 1F**). Within the cortex, most neuron types were broadly distributed across subregions, yet many displayed pronounced regional enrichment or depletion. Notably, several neuron types exhibited spatial gradients across cortical areas. For example, consistent with classical cytoarchitecture(*17*), the L4 IT neurons were absent in agranular insular subregions (AId, AIv, and AIp) but became progressively enriched across dysgranular and granular regions (gustatory cortex, GU, also known as dysgranular IC; visceral cortex, VISC, also known as granular IC; somatosensory cortex, SSC) (**Fig. 1F**). Conversely, the L2/3 IT 1 and L6 CT 2 neurons were enriched in the agranular IC but depleted in granular IC and SSC, whereas L2/3 IT 2 and L6 CT 1 displayed the inverse pattern (**Fig. 1F**). Spatial mapping of individual subtypes onto representative coronal sections resolved highly precise laminar and subregional localizations (**Fig. 1G and fig. S4C**). These discrete enrichments and continuous gradients define a sharp cellular boundary between the IC and the ventrally adjacent Pir, contrasted by a gradual transition toward dorsal neocortical areas, which may underlie the distinct connectivity and functional specialization of these subregions.

Given the distinct functions and cellular organization of the IC and adjacent areas, we next examined whether these regions exhibit specialized transcriptional features. Comparative analysis identified 61 genes significantly enriched and 54 depleted in the IC compared to adjacent cortical areas within our MERFISH library (**fig. S5A**) (**table S5**). Spatial mapping of representative enriched (*Lypd1*, *Smoc2*) and depleted (*Lmo3*, *Scn4b*) genes onto the anterior and posterior coronal sections confirmed their distinct distribution patterns (**fig. S5B**). These profiles were highly consistent with *in situ* hybridization data from Allen Brain Atlas (**fig. S5C**), validating our MERFISH measurements. To extend our spatial analysis to a whole-transcriptome scale, we integrated our MERFISH data with a public IC scRNA-seq dataset(*30*) to impute genome-wide spatial expression profiles(*32*) (Methods). This whole-transcriptome imputation identified 2,444 enriched and 1,153 depleted genes in the IC compared to adjacent cortical areas (**fig. S5D**) (**table S5**). Gene Ontology enrichment analysis of these genes revealed that IC enriched genes were strongly associated with multiple metabolic processes, whereas IC depleted genes were associated with myelination and axon ensheathment (**fig. S5E**). Finally, we evaluated whole-transcriptome expression patterns across specific anatomical subregions, identifying regional marker genes such as *Scn5a* in the Pir, *Nnat* in the EPd and CLA, and *Krt77* in the AIp (**fig. S5F**). Together, these results demonstrate that the IC subregions and adjacent regions have a discrete molecular architecture, which may underlie their specialized neuronal organization and functional properties.

Collectively, the profiling and analysis establish a high-resolution spatial single-cell map of the IC and adjacent regions, providing a critical foundation for deciphering the neural connectivity and functional architecture of the IC.

### Projection patterns of transcriptomically defined neuron types in IC follow a multiple-to-multiple architecture

To investigate the relationship between transcriptomic identity and projection patterns of the IC neurons, we integrated MERFISH with retrograde tracing to simultaneously decode the spatial transcriptomes and long-range projections(*16*) (Methods). To this end, we injected cholera toxin subunit B (CTB) conjugated to spectrally distinct fluorophores into the major IC projection targets to retrogradely label the projecting neurons in IC. Serial coronal sections containing the IC and adjacent regions were then processed for CTB imaging to identify projection-specific neurons, followed by MERFISH to determine their transcriptomic profiles (**Fig. 2A**). Spatial maps were subsequently registered to the Allen CCFv3 for anatomical annotation and 3D reconstruction (**Fig. 2A**).

**Fig. 2.**
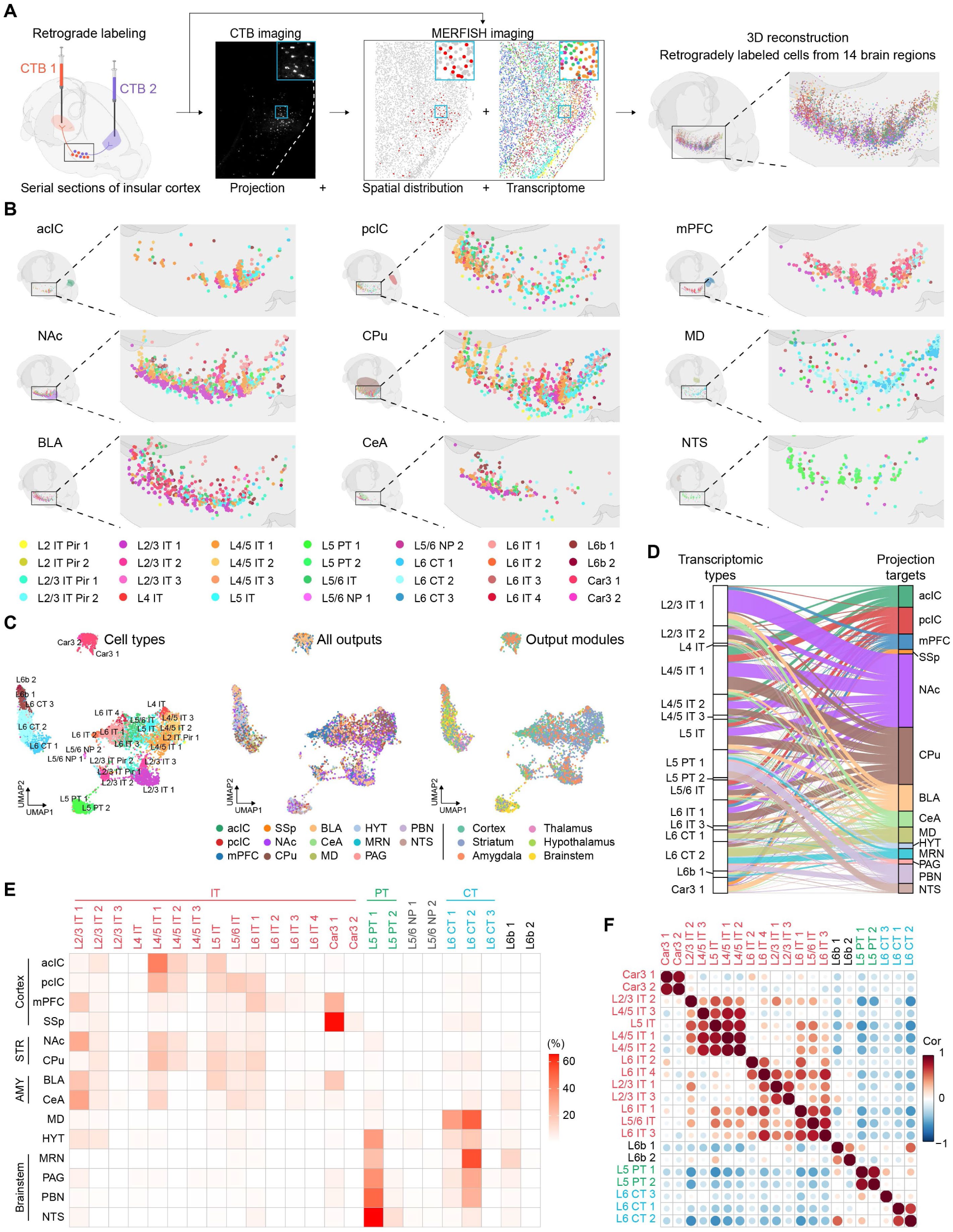
Comprehensive mapping of spatially resolved transcriptomic identities and projection patterns. **(A)** Workflow that integrates MERFISH with retrograde tracing to determine the transcriptomic identities of the projecting neurons. Shown are representative images of CTB-labeled cells in the IC and MERFISH results decoding the spatial transcriptomics of the cells, as well as 3D reconstruction of all retrogradely labeled cells in the IC (from 14 major downstream targets in 7 mice after registration to Allen CCFv3). **(B)** 3D maps of the retrogradely labeled neurons in the IC with transcriptomic identity for individual downstream targets. The cells are color-coded by their transcriptomic identities. **(C)** UMAP visualization of all retrogradely labeled cells identified by MERFISH. The cells are color-coded by their transcriptomic identities (left), the exact projection targets (middle) and categories of projection targets (right). **(D)** The projection targets of molecularly defined excitatory neuron subtypes in the IC, represented by an alluvial diagram. Each line represents a labeled cell, and the colors indicate the individual projection targets. **(E)** Heatmap showing the proportion of the transcriptomic types in the IC for the specific projection targets. The IT, PT and CT subtypes are labeled in red, green and blue, respectively. STR, striatum; AMY, amygdala. **(F)** Heatmap showing the correlation of the projection similarity between the transcriptomic types in the IC. The NP neuron subtypes were not included in this analysis due to limited cell numbers.

To comprehensively profile the IC output organization, we mapped the projections to 14 major brain regions previously identified(*16*) across 7 mice. These included two contralateral cortical targets, the anterior IC (acIC) and posterior IC (pcIC), and twelve ipsilateral cortical and subcortical targets: medial prefrontal cortex (mPFC), primary somatosensory cortex (SSp), nucleus accumbens (NAc), caudate putamen (CPu), basolateral amygdala (BLA), central amygdala (CeA), mediodorsal thalamus (MD), hypothalamus (HYT), midbrain reticular nucleus (MRN), periaqueductal grey (PAG), parabrachial nucleus (PBN), and nucleus of the solitary tract (NTS). Following validation of each injection site (**fig. S6A-N**), retrogradely labeled neurons in the IC and adjacent regions were processed to reconstruct the 3D spatial distribution alongside transcriptomic identity (**Fig. 2B**). As an additional specificity control, we injected CTB into the spinal vestibular nucleus (SpVe), a neighboring region to NTS that does not receive projections from the IC(*16*), and we detected no retrograde labeling in the IC (**fig. S6O**), supporting the specificity of our approach. Overall, we identified 6,227 retrogradely labeled neurons in the IC and adjacent regions across 14 major projection targets (**table S4**).

Mapping individual projection targets onto the transcriptomic space showed no discrete, target-specific clustering, although certain targets displayed enrichment within specific transcriptomic types (**Fig. 2C**). To clarify these distributions, we grouped the 14 projection targets into six broader anatomical categories. These spanned intratelencephalic targets (cortices, striatum, and amygdala) and extratelencephalic targets (thalamus, hypothalamus, and brainstem). We then projected these categories either collectively (**Fig. 2C, right**) or individually (**fig. S6P**) onto the transcriptomic space. Neurons targeting cortices, striatum and amygdala mapped predominantly to transcriptomic IT classes. Conversely, neurons projecting to thalamus, hypothalamus and brainstem mainly belonged to PT and CT classes. Notably, projections to the MD originated almost exclusively from the CT neurons (**Fig. 2C and fig. S6P**).

To further examine the relationship between transcriptomic identity and projection patterns, we directly cross-mapped these two features (**Fig. 2D**) and quantified the cell-type composition for each projection target (**Fig. 2E**). We found that individual transcriptomic types innervate multiple projection targets, and conversely, most projection targets receive innervation from multiple transcriptomic types, demonstrating a multiple-to-multiple architecture of correspondence (**Fig. 2, D and E**). Notably, distinct projection motifs emerged across the major neuron classes. The IT neurons preferentially innervate the intratelencephalic structures, including cortices, striatum and amygdala. The PT neurons project to the hypothalamus and specific brainstem nuclei, including the MRN, PAG, PBN, and NTS, whereas the CT neurons strongly target the thalamus while also extending projections to the brainstem (**Fig. 2, D and E**).

Despite the absence of clear one-to-one mapping, individual transcriptomic types exhibited distinct projection preferences. For instance, while the projections to cortical areas originate from multiple IT neurons, the contralateral IC receives more projections from deep-layer IT neurons, whereas the SSp is preferentially innervated by the Car3 population (**Fig. 2, D and E**). While the NAc and CPu both received robust IT inputs with highly similar cell type compositions, the L2/3 IT 1 neurons showed much greater enrichment projecting to the NAc (**Fig. 2, D and E**). Interestingly, the BLA and CeA displayed distinct innervation profiles, with L5 IT and Car3 1 neurons preferentially targeting the BLA whereas L2/3 IT 1 and L4/5 IT 1 preferentially targeting the CeA (**Fig. 2, D and E**). Moreover, some downstream targets also exhibit clear cell type preferences. For instance, the MD receives nearly exclusive inputs from CT subtypes, whereas the HYT integrated inputs from IT, PT, and CT neurons (**Fig. 2, D and E**). Notably, and contrary to classical models depicting CT neurons as exclusively thalamic projecting, both PT and CT neurons projected to brainstem. Within the brainstem, CT neurons preferentially innervate the MRN, whereas PT neurons primarily target the NTS (**Fig. 2, D and E**). Finally, quantitative comparison of connectivity matrices demonstrated that projection patterns are highly conserved within major neuron classes but diverge sharply between them (**Fig. 2F**), indicating the IT, PT, and CT neuron classes as discrete neural substrates with distinct transcriptomic and projection profiles.

Collectively, these results demonstrate that IC neuron projection patterns follow a multiple-to-multiple architecture characterized by distinct cell-type biases: each transcriptomically distinct neuron type projects to multiple downstream targets, and each downstream target receives projections from multiple transcriptomic types. Major neuron classes exhibit segregated projection profiles. IT neurons predominantly target cortical regions, striatum and amygdala. Conversely, PT neurons project preferentially to the hypothalamus and brainstem, whereas CT neurons innervate the thalamus while also extending to specific brainstem nuclei.

### Transcriptomic identity and spatial location jointly govern neuronal projection patterns

To study the relationship among transcriptomic identity, spatial location and projection targets of the neurons, we mapped the spatial distribution of the single neurons retrogradely labeled from specific targets (**Fig. 3A**). The coronal slices from different mice were from similar locations along the AP axis and registered to Allen CCFv3 to ensure proper anatomical annotation. Distinct topographic distributions of the retrogradely labeled neurons were revealed along the AP axis and across anatomical subregions (**Fig. 3A and fig. S7, A and B**). Along the AP axis, neurons projecting to the acIC, CPu, or MD predominantly localized to anterior sections (**Fig. 3A and fig. S7A**). Conversely, projections to pcIC or CeA preferentially originate from posterior sections, and neurons targeting the mPFC are concentrated within intermediate sections, whereas the remaining downstream targets show no pronounced AP bias (**Fig. 3A and fig. S7A**). Subregional mapping further resolved distinct anatomical biases (**fig. S7B**). For example, neurons projecting to the acIC primarily arise from the anterior agranular IC (AId and AIv). In contrast, pcIC-projecting neurons are distributed broadly across the entire IC but absent from the CLA. For striatal targets, the NAc receives inputs mainly from the agranular IC, whereas the CPu is innervated broadly from the entire IC and biased towards the AId (**Fig. 3A and fig. S7B**). Together, these findings reveal that the projection neurons are highly organized spatially in the IC with distinct topographic distribution features.

**Fig. 3.**
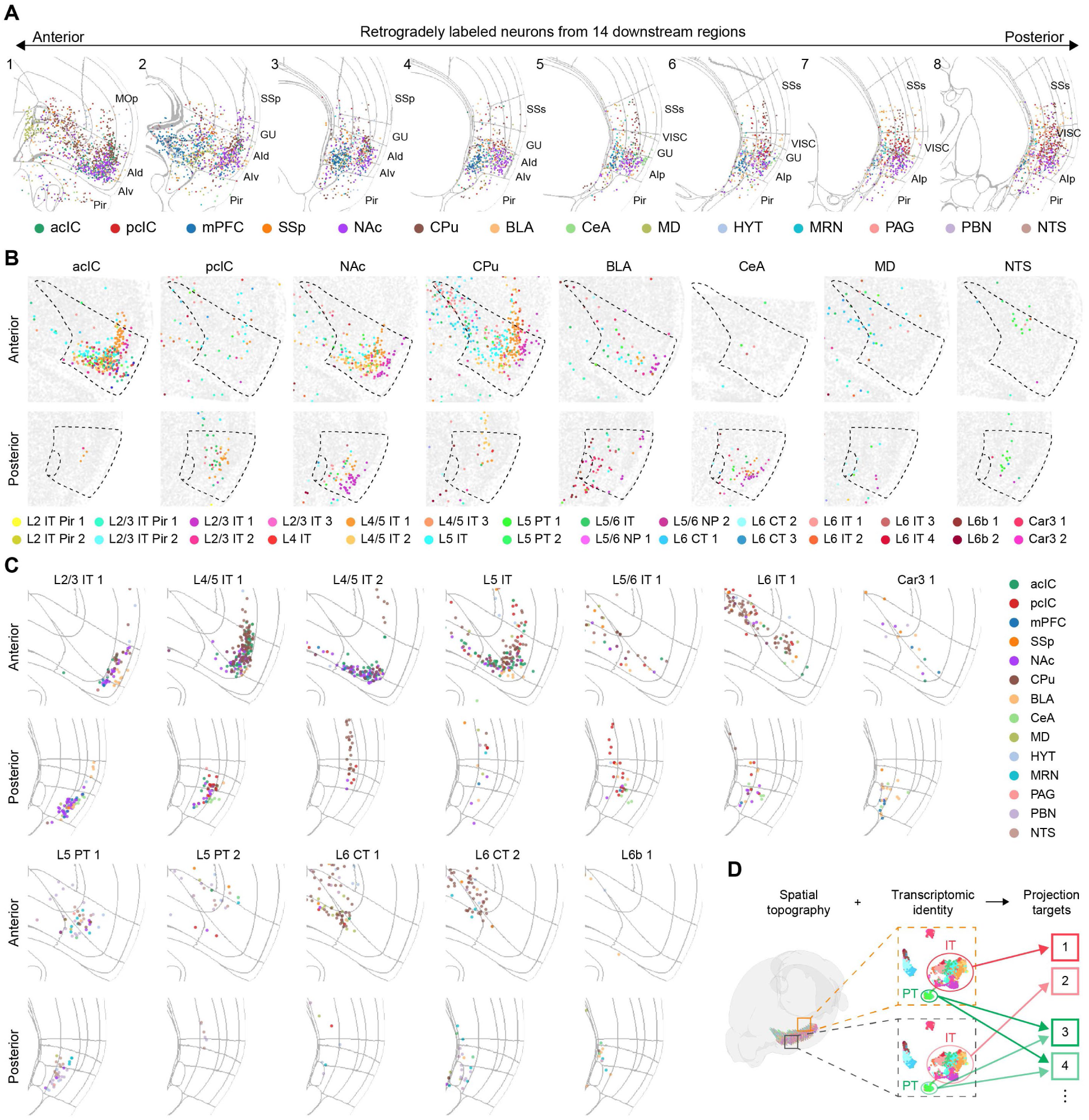
Topographically varied projection patterns of transcriptomic neuron types. **(A)** Reconstructed overlaid images of all retrogradely labeled cells from 14 projection targets in the serial coronal slices spanning from the anterior to posterior sections of the IC and adjacent regions. The neurons were plotted to the representative Allen CCFv3 sections for display and color-coded by the projection targets. **(B)** Representative spatial maps of the anterior (top) and posterior (bottom) coronal slices containing the IC showing the topographic distribution of the retrogradely labeled cells for each individual projection target. The cells are color-coded by their transcriptomic identities. **(C)** Representative maps of the anterior and posterior IC showing the topographic projection patterns for individual excitatory neuron subtypes. The cells are color-coded by the projection targets. **(D)** Schematic diagram of the spatially dependent projection patterns of transcriptomic types. The same transcriptomic types with different spatial distributions have similar (PT neurons) or distinct (IT neurons) projection targets.

To determine how transcriptomic identity and projection pattern are spatially correlated, we mapped the transcriptomic identities and topographic distribution of retrogradely labeled neurons in 2D coronal sections (**Fig. 3B and fig. S7, C and D**). Individual downstream targets received projections from multiple transcriptomic neuron types that exhibited highly stereotyped spatial patterns in the IC and adjacent regions (**Fig. 3B and fig. S7D**). Contralateral projections to the acIC primarily arise from layer 2 to 5 IT neurons in the anterior IC (aIC), with a distinct ventral enrichment in L5 IT neurons and notably, few labeled neurons were detected in the posterior IC (pIC). In contrast, projections to pcIC originated from both the aIC and pIC but exhibit diverging cellular compositions: the projections are dominated by L5 IT neurons in aIC, whereas in pIC the neurons are enriched with L4/5 IT 1, L4/5 IT 2, and L5/6 IT subtypes (**Fig. 3B**). Striatal projections also show distinct spatial and transcriptomic segregation between the NAc and CPu (**Fig. 3B**). For the NAc projections, L2/3 IT neuron numbers are comparable in aIC and pIC, while the L4/5 IT and L5 IT neurons are markedly depleted in the pIC compared to the aIC. For the CPu projections, much fewer neurons are detected in the pIC, with L4/5 IT 2 neurons as the major transcriptomic type. There are higher abundances of L2/3 IT, L4/5 IT, L5 IT and L6 IT neurons in the aIC, indicating different transcriptomic types in different subregions of IC projecting to the CPu. Notably, while neurons targeting the NAc and CPu are largely intermingled in the aIC, they are spatially segregated in the pIC (**Fig. 3B**). Similarly, the BLA receives projections from distinct transcriptomic types in aIC and pIC despite the L2/3 IT neurons are labeled in both aIC and pIC (**Fig. 3B**). Interestingly, the NTS is innervated by the L5 PT neurons uniformly across the subregions (**Fig. 3B**). Together, these results demonstrate that downstream targets receive projections from heterogeneous transcriptomic neuron types, which are spatially organized into specific subregional and laminar configurations.

To further determine how the correspondence between transcriptomic identity and connectivity varies spatially, we examined the projection profiles of individual transcriptomic neuron types. Each neuron type projects to multiple downstream targets, with the projection patterns shifting across anatomical subregions (**Fig. 3C**). First, projection patterns for most transcriptomic types varied along the AP axis. For example, while the L4/5 IT 1 neurons target the NAc and BLA in both aIC and pIC, their other targets inverted along the AP axis: the L4/5 IT 1 neurons in aIC robustly innervate the acIC and CPu but not the pcIC and CeA, whereas in the pIC the same transcriptomic type targets pcIC and CeA but not the acIC and CPu (**Fig. 3C**). Similarly, the L5 IT neurons in aIC project strongly to the acIC, pcIC, NAc, CPu, and BLA, whereas in the pIC they exhibit minimal projections to these targets (**Fig. 3C**). In contrast to these highly dynamic IT projections, L5 PT 1 neurons maintain stable projection profiles along the AP axis, consistently targeting multiple brainstem nuclei including the PAG, PBN, and NTS (**Fig. 3C**). Second, projection patterns for most IT neurons shift continuously along the dorsoventral or mediolateral axes. For instance, within the aIC, L4/5 IT 1 and L4/5 IT 2 neurons projecting to the NAc are segregated ventrally, whereas those projecting to the CPu are localized dorsally, with cells situated at intermediate positions projecting to both targets (**Fig. 3C**). This topographical logic is conserved for the L5 IT neurons in aIC, which further display a ventral bias for BLA projections and a dorsal bias for pcIC projections. Conversely, the L5 PT 1 neurons projecting to different targets in brainstem are largely intermingled, lacking clear spatial segregation (**Fig. 3C**). Interestingly, challenging the classical view that CT neurons exclusively target the thalamus, we found that the L6 CT 2 neurons situated in layer 6b or surrounding the CLA project to the MRN and PBN (**Fig. 3C**), indicating that the connectivity of this specialized L6b CT subpopulation diverges from the canonical layer 6 CT neurons. Together, these results demonstrate that each transcriptomic type possesses highly structured, spatially dependent projection patterns.

Collectively, our findings suggest that the neuronal projection patterns are specified by transcriptomic identity and spatial topography (**Fig. 3D**). The major neuron classes (IT, PT, and CT neurons) innervate distinct sets of downstream targets, while the transcriptomic types within the classes target overlapping regions with distinct spatially dependent patterns.

### Impacts of chronic neuropathic pain on the IC neurons

Given the critical roles of the IC in pain processing, we next investigated how chronic neuropathic pain impacts neurons with distinct transcriptomic and projection profiles. To this end, we performed spared nerve injury (SNI) to induce chronic neuropathic pain, with sham surgery serving as a control (**fig. S8A**). A significant decrease in mechanical withdrawal thresholds in SNI mice was revealed by von Frey test (VFT) (**fig. S8B**), confirming the tactile allodynia in the mice. Four weeks after the surgery, brain sections encompassing the IC and adjacent regions were then collected for MERFISH analysis. Unsupervised clustering analysis indicated no broad segregation between the sham and SNI groups with few differentially expressed genes (DEGs) detected (**fig. S8C**), indicating global transcriptomes are not altered in chronic neuropathic pain. To identify the neuron populations potentially perturbed by chronic pain, we employed Augur analysis(*33*) and identified L5 PT 2 neurons as the most perturbed transcriptomic type, followed by L6 CT 3 and L4/5 IT 3 (**fig. S8D**). Since the IC is activated in chronic pain(*21*), we examined whether these changes could be detected through the baseline expression of activity-related genes (ARGs). We calculated an aggregate ARG score based on the mean expression of a panel of five ARGs (Methods) and found L5 PT 2 as the only transcriptomic type showing a significant increase in ARG score (**fig. S8E**). Collectively, these results indicate that chronic neuropathic pain drives transcriptomic perturbations across multiple cell types with elevated baseline activity within specific neuronal subpopulations.

### Discrete neural networks of molecularly defined neuron types

We next investigated whether molecularly defined neuron populations in the IC play different roles in pain processing. Given the distinct projection patterns of the IT, PT and CT neurons, and the transcriptomic perturbations of chronic pain on multiple cell types as described above, we hypothesized that these neuronal populations are differentially involved and play different roles in pain through the distinct projections. Thus, we targeted these three molecularly defined neuron types for functional dissection: the L5 IT neurons (marked by the expression of *Il1rapl2*) as a representative population of IT neurons, the PT neurons (marked by *Fezf2*), and the CT neurons (marked by *Foxp2*) (**fig. S1C and S9A**). For the remainder of the study, we focused on the pIC because it plays a critical role in multimodal integration of sensory information(*19, 34–36*) and regulates different dimensions of pain(*18, 26, 27*). Furthermore, its compact anatomical volume encompasses the diverse transcriptomic types of IC neurons, facilitating specific and localized viral tracing, functional imaging and chemogenetic manipulations.

To determine how these molecularly distinct neuron types integrate into the pain processing network, we mapped their brain-wide projections. To this end, we injected a Cre-dependent adeno-associated virus (AAV) expressing mGFP and presynaptic marker synaptophysin-fused mRuby into the pIC of *Il1rapl2*-Cre, *Fezf2*-CreER, or *Foxp2*-Cre mice (**Fig. 4A**). The virus expression was confirmed by *post hoc* histological examination (**fig. S9B**). Consistent with the correspondence between transcriptomic identity and projection pattern, brain-wide mapping revealed distinct projection profiles for the IT^Il1rapl2^, PT^Fezf2^ and CT^Foxp2^ neurons in pIC (**Fig. 4B and fig. S9C-F**). The IT^Il1rapl2^ neurons project to the lateral NAc, ventral CPu, dorsal peduncular cortex (DP), BLA, and CeA, with no detectable signals in the thalamus or brainstem. In contrast, the PT^Fezf2^ neurons project broadly to the thalamus, PAG, PBN, rostral ventromedial medulla (RVM) and NTS, alongside minor projections to the BLA, CeA and striatum, whereas the CT^Foxp2^ neurons predominantly target the thalamus with only very weak signals in the PAG. Quantification of the mRuby fluorescence confirmed these highly segregated projection patterns (**Fig. 4C**). Interestingly, the IT^Il1rapl2^ and PT^Fezf2^ neurons project to the amygdala with distinct subregional topography, with the IT^Il1rapl2^ neurons strongly innervating the BLA and capsular CeA (CEAc), whereas the PT^Fezf2^ preferentially targeting the medial CeA (CEAm) (**fig. S9C**). Together, these results demonstrate the divergent outputs of the IT^Il1rapl2^, PT^Fezf2^ and CT^Foxp2^ neurons in pIC, which potentially underlies their different functions in pain processing.

**Fig. 4.**
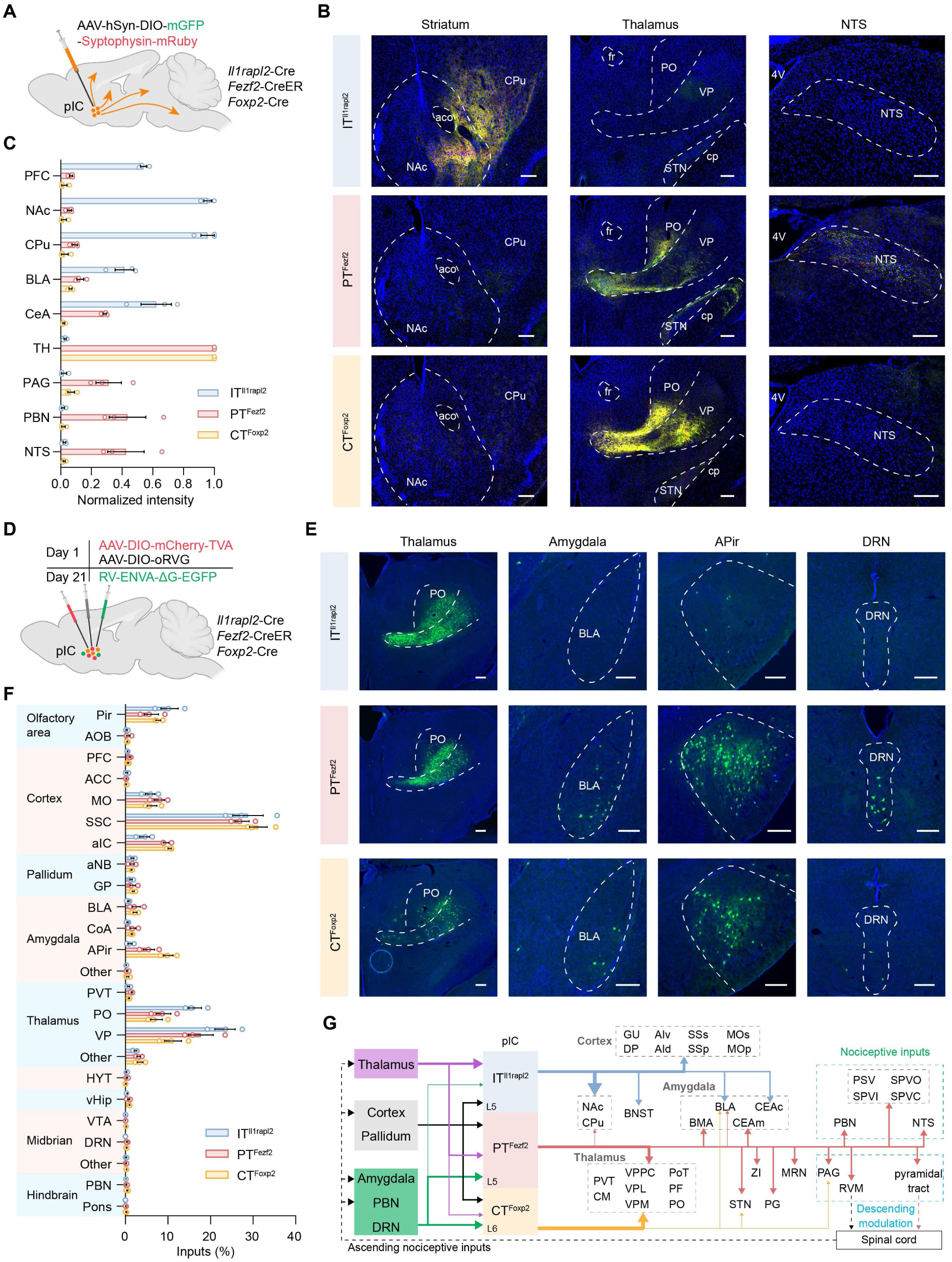
Molecularly defined neuron types in the pIC are wired into discrete neural networks. **(A)** Diagram of anterograde tracing for the IT^Il1rapl2^, PT^Fezf2^ and CT^Foxp2^ neurons in pIC. **(B)** Representative images showing the projections of the IT^Il1rapl2^, PT^Fezf2^ and CT^Foxp2^ neurons to the striatum (left), thalamus (middle) and NTS (right). Scale bar, 200 μm. **(C)** Quantification of the normalized average mRuby fluorescence intensity to the major projection targets. n = 3 mice. **(D)** Diagram of monosynaptic retrograde rabies tracing of the IT^Il1rapl2^, PT^Fezf2^ and CT^Foxp2^ neurons in pIC. **(E)** Representative images of the retrogradely labeled monosynaptic input neurons in the thalamus, amygdala, APir and DRN. Scale bar, 200 μm. **(F)** Quantification of the monosynaptic input neurons across the brain showing the fractions of the total input cells. n = 3 mice. **(G)** Summary wiring diagram of efferent and afferent connections of the IT^Il1rapl2^, PT^Fezf2^ and CT^Foxp2^ neurons in IC. The strokes of the lines indicate the strength of the projections, and the dashed lines indicate the known innervation based on the literature. Abbreviations are in table S6.

We next examined the monosynaptic inputs to the IT^Il1rapl2^, PT^Fezf2^ and CT^Foxp2^ neurons using rabies virus (RV) retrograde tracing. To this end, Cre-dependent AAV helpers encoding TVA-mCherry and rabies G protein were injected into the pIC of *Il1rapl2*-Cre, *Fezf2*-CreER, or *Foxp2*-Cre mice, followed by EnvA-ΔG rabies virus expressing EGFP injected into the same region three weeks later (**Fig. 4D**). One week after the RV injection, the virus expression and starter cells were confirmed in the injection sites (**fig. S10A**). Brain-wide examination of the RV-EGFP labeled neurons revealed that the pIC IT^Il1rapl2^, PT^Fezf2^ and CT^Foxp2^ neurons received major monosynaptic inputs from multiple brain regions, including the somatosensory cortex (SSC), motor cortex (MO), aIC, Pir, posterior (PO) and ventroposterior (VP) complexes of thalamus (**Fig. 4, E and F and fig. S10B-E**). Interestingly, while the pIC IT^Il1rapl2^, PT^Fezf2^ and CT^Foxp2^ neurons receive comparable monosynaptic inputs from the pallidum and cortices such as MO and SSC, their subcortical inputs diverged sharply (**fig. S10B-E**). Specifically, the IT^Il1rapl2^ neurons integrate a higher proportion of inputs from the PO and VP thalamus, but fewer inputs from the BLA, amygdalopiriform transition area (APir), and dorsal raphe nucleus (DRN) compared to the Fezf2^CreER^ or Foxp2^Cre^ neurons (**Fig. 4E**). The different upstream connectivity patterns indicate that these neuron classes are differentially positioned in the circuits and may therefore process distinct incoming nociceptive information.

Collectively, these results reveal that the pIC IT^Il1rapl2^, PT^Fezf2^ and CT^Foxp2^ neurons are differentially integrated into the neural networks (**Fig. 4G**). Each neuron class may integrate a unique composition of nociceptive information via the monosynaptic inputs and coordinate distinct downstream functional outputs during pain processing.

### Differential recruitment of molecularly defined neuron types in pain processing

To explore the potential function of the pIC IT^Il1rapl2^, PT^Fezf2^ and CT^Foxp2^ neurons in pain processing, we performed *in vivo* fiber photometry to record the neuronal activity. To this end, we injected Cre-dependent AAV expressing jGCaMP7s, a Ca^2+^ indicator, into the pIC of *Il1rapl2*-Cre, *Fezf2*-CreER, or *Foxp2*-Cre mice, and implanted an optic fiber above the virus injection site (**Fig. 5A**). Histological examination confirmed the virus expression and fiber placement (**Fig. 5B**). Three weeks after the surgery, the Ca^2+^ dynamics were recorded to indicate the neuronal activity during noxious or innocuous stimulations (**Fig. 5C**).

**Fig. 5.**
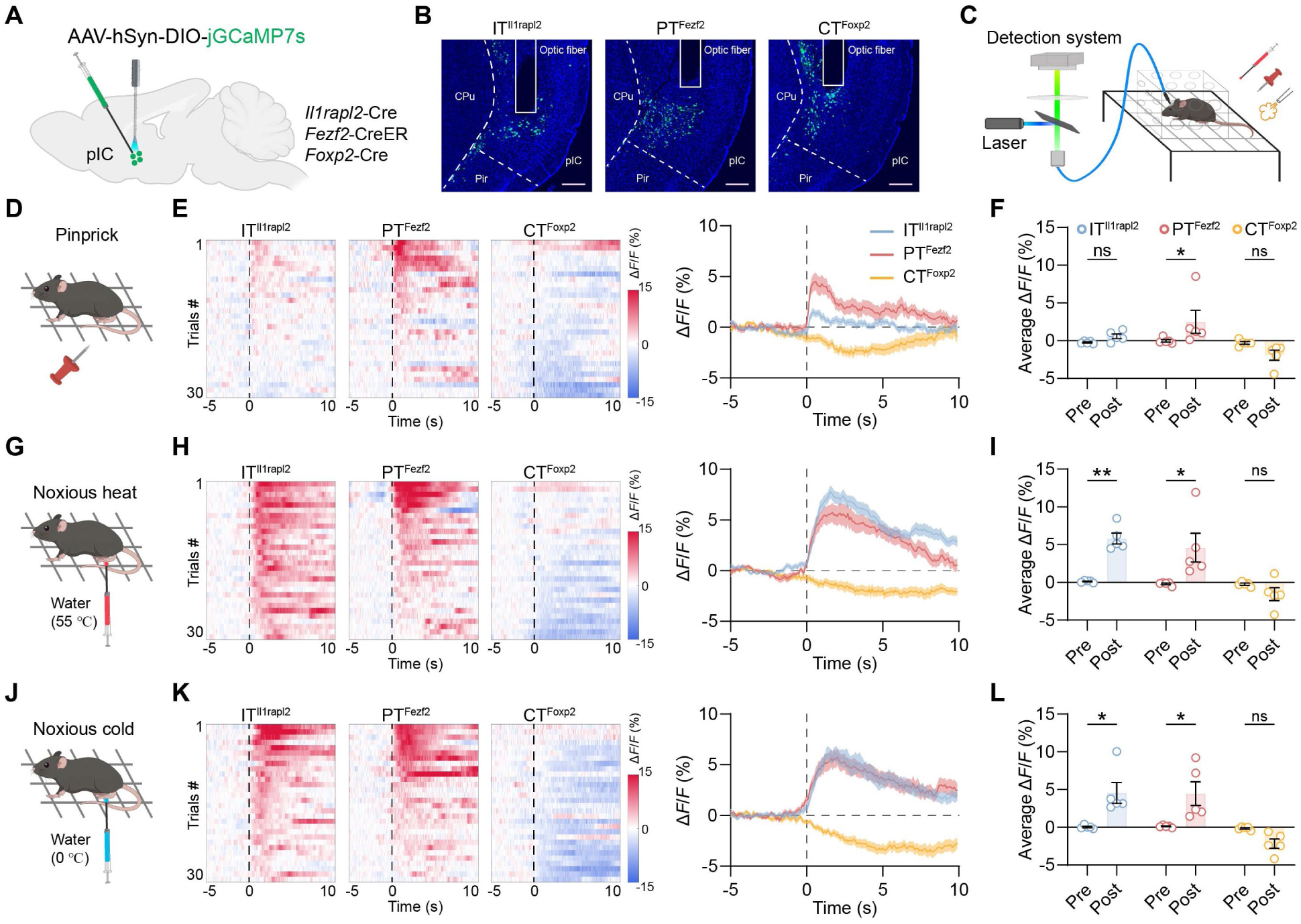
Molecularly defined neuron types in the pIC differentially respond to noxious stimulation. **(A)** Schematic diagram of virus injection and optical fiber implantation into the pIC of the *Il1rapl2*-Cre, *Fezf2*-CreER or *Foxp2*-Cre mice. **(B)** Representative images showing the virus expression and fiber implantation in the pIC. **(C)** Diagram of the optical fiber photometry assay of the mice subjected to various stimuli. **(D)** Diagram of pinprick stimulation to the mice. **(E)** Heatmaps (left) and averaged traces (right) showing the calcium activity of the IT^Il1rapl2^ (blue), PT^Fezf2^ (red) or CT^Foxp2^ (yellow) neurons in the pIC during pinprick stimulation. The dashed lines indicate the onset of pinprick stimulation. The solid lines and shadow indicate the mean and s.e.m., respectively. n = 5 mice and 6 trials for each mouse. **(F)** Averaged ΔF/F (%) of the three neuron populations in the pIC 5 s before (pre) and after (post) the stimulation. RM two-way ANOVA with Sidak’s multiple comparison tests. **(G-I)**, similar to **(D-F)** but for noxious heat stimulation with hot water (55 °C). **(J-L)**, similar to **(D-F)** but for noxious cold stimulation with cold water (0 °C). \**P* < 0.05; \*\**P* < 0.01; ns, not significant.

To assess the involvement of the IT^Il1rapl2^, PT^Fezf2^ and CT^Foxp2^ neurons in mechanical pain processing, we subjected the mice to pinprick stimulation and recorded the Ca^2+^ dynamics (**Fig. 5D**). The PT^Fezf2^ neurons exhibited a robust increase in Ca^2+^ signals compared to the baseline, whereas the IT^Il1rapl2^ neurons showed a minor increase and the CT^Foxp2^ neurons displayed a decrease in Ca^2+^ signals (**Fig. 5E**). Quantification confirmed the significant increase in Ca^2+^ signals specifically in the PT^Fezf2^ neurons, with no significant changes in the IT^Il1rapl2^ or CT^Foxp2^ neurons (**Fig. 5F**), indicating specific activation of the PT^Fezf2^, but not the IT^Il1rapl2^ or CT^Foxp2^ neurons, in response to noxious mechanical stimulation. The mice were also subjected to innocuous light touch stimulation, and no significant changes in the Ca^2+^ signals were observed in the PT^Fezf2^, IT^Il1rapl2^ or CT^Foxp2^ neurons (**fig. S11A-C**), indicating that these neuron populations do not strongly respond to innocuous mechanical stimulation. Together, these results indicate that the PT^Fezf2^ neurons are specifically activated in mechanical pain processing.

We next evaluated the involvement of these neuron populations in thermal pain processing by applying noxious heat (**Fig. 5G**). Both the IT^Il1rapl2^ and PT^Fezf2^ neurons showed robust increase in Ca^2+^ signals in response to noxious heat stimulation, whereas the CT^Foxp2^ neurons exhibited a decreasing trend (**Fig. 5H**). Quantitative analysis confirmed the significant increase in Ca^2+^ signals in both the PT^Fezf2^ and the IT^Il1rapl2^ neurons (**Fig. 5I**), indicating that these two populations are significantly activated in response to noxious heat stimulation. Similarly, noxious cold stimulation elicited significant increase in Ca^2+^ signals in both the IT^Il1rapl2^ and PT^Fezf2^ neurons and again, a decrease in the CT^Foxp2^ neurons (**Fig. 5J-L**). The mice were also subjected to innocuous warmth stimulation, and a minor but significant increase in Ca^2+^ signals was observed in the PT^Fezf2^ neurons (**fig. S11D-F**). A minor increase in the Ca^2+^ signals of IT^Il1rapl2^ neurons was also observed, while no dramatic change was detected in the CT^Foxp2^ neurons (**fig. S11D-F**). Together, these results indicate that both the IT^Il1rapl2^ and PT^Fezf2^ neurons are significantly activated in response to noxious thermal stimulation, whereas the CT^Foxp2^ neurons might be inhibited. Lastly, we tested whether these neuron populations respond to innocuous aversive stimuli using an air puff assay (**fig. S11G**). We found a sharp, although statistically not significant, increase in the Ca^2+^ signals in the IT^Il1rapl2^ and PT^Fezf2^, but not the CT^Foxp2^ neurons, in response to air puff stimulation (**fig. S11, H and I**), indicating that in the IT^Il1rapl2^ and PT^Fezf2^, but not the CT^Foxp2^ neurons, might also involve in the processing of innocuous aversive stimulation.

In summary, these results demonstrate that the PT^Fezf2^ neurons are activated in response to both noxious mechanical and thermal stimulation, and that the IT^Il1rapl2^ neurons are specifically activated in response to noxious thermal but not mechanical stimulation, while the CT^Foxp2^ neurons showed a trend of decreased activity in response to both mechanical and thermal stimulation (**fig. S11J**). These results reveal that these molecularly defined neuron types are differentially involved in different dimensions of pain, indicating potential segregated pain processing pathways within the pIC.

### Distinct functions of molecularly defined neuron types in pain processing

Given the distinct neural networks and neuronal activity of the IT^Il1rapl2^, PT^Fezf2^ and CT^Foxp2^ neurons in response to noxious stimulation, we next asked whether these neuron populations play different roles in pain regulation. To this end, Cre-dependent AAVs expressing hM4D(Gi), a chemogenetic inactivator, were bilaterally injected into the pIC of *Il1rapl2*-Cre, *Fezf2*-CreER, or *Foxp2*-Cre mice with AAVs expressing mCherry as a control (**Fig. 6A**). Three weeks later, behavioral assays were performed 20 min after an intraperitoneal injection of clozapine-N-oxide (CNO). Virus expression was confirmed by *post hoc* histological examination (**Fig. 6B**).

**Fig. 6.**
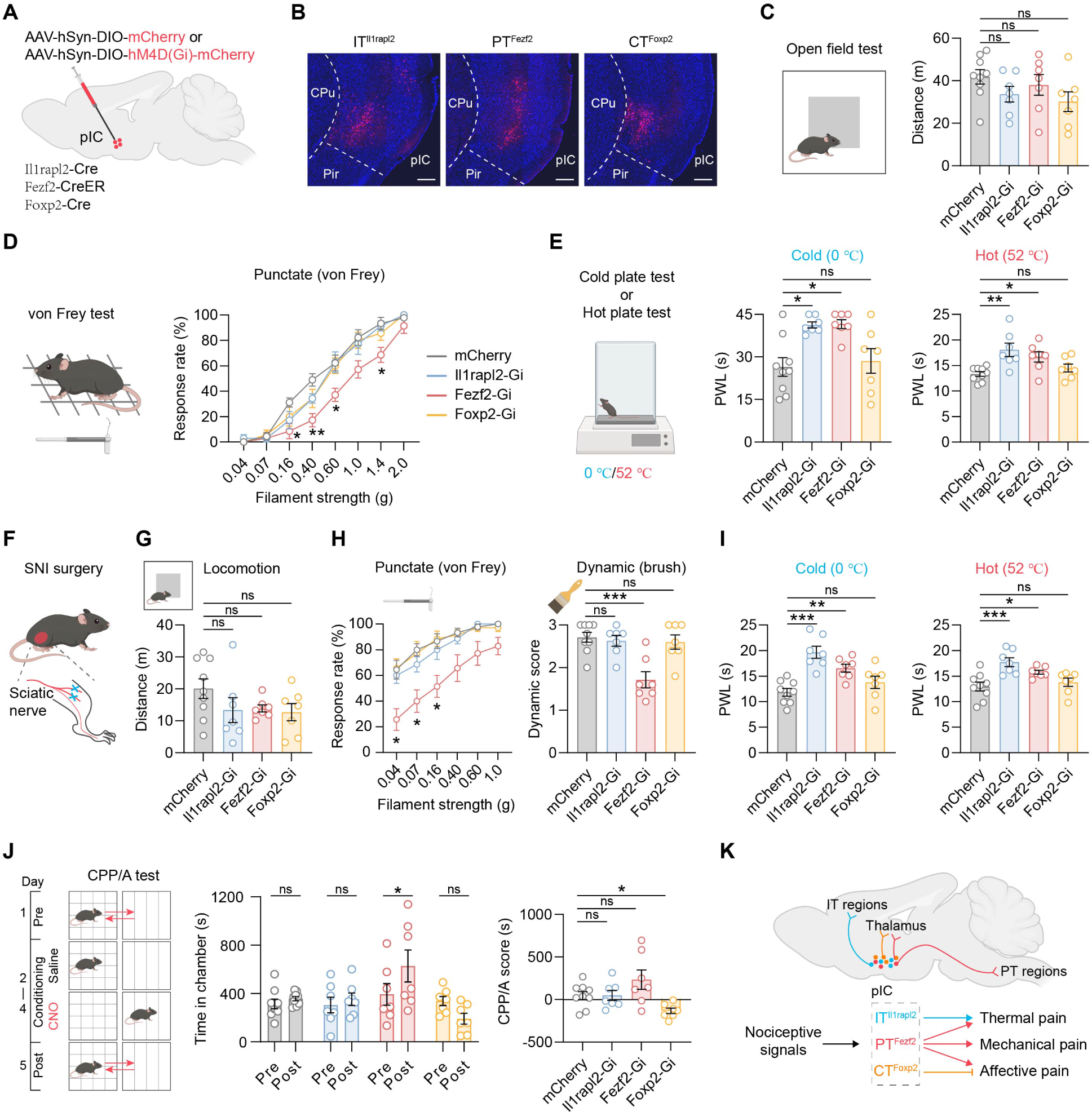
Different functions of the molecularly defined neuron types in pIC in pain processing. **(A)** Diagram of virus injection into the pIC of the *Il1rapl2*-Cre, *Fezf2*-CreER or *Foxp2*-Cre mice. **(B)** Representative images confirming the virus expression in the pIC. **(C)** Diagram of open field test (left), and the traveled distance of the mice of mCherry (grey; n = 9 mice), Il1rapl2-Gi (blue; n = 7 mice), Fezf2-Gi (red; n = 7 mice) and Foxp2-Gi (yellow; n = 7 mice) groups. The same numbers of animals were used for all the experiments below. One-way ANOVA with Dunnett’s multiple comparison tests. **(D)** Diagram of von Frey test (left), and the positive response rates to the von Frey filament stimulation (right) of the mice. RM two-way ANOVA with Tukey’s multiple comparison tests. **(E)** Diagram of hot/cold plate test (left), and the paw withdrawal latency (PWL) of the mice subjected to the cold plate (middle) or hot plate (right). Kruskal-Wallis test with Dunn’s multiple comparison tests for cold plate test and one-way ANOVA with Dunnett’s multiple comparison tests for hot plate test. **(F)** Diagram of SNI surgery to induce chronic neuropathic pain. **(G)** Traveled distance of the mice in open field test 4 weeks after SNI surgery. Brown-Forsythe and Welch ANOVA test with Dunnett’s T3 multiple comparison test. **(H)** Mechanical punctate or dynamic allodynia of the mice after SNI surgery as measured in von Frey test (left) or brush test (right). RM two-way ANOVA with Tukey’s multiple comparison tests for von Fret test and one-way ANOVA with Dunnett’s multiple comparison tests for brush test. **(I)** Cold (left) and heat (right) sensitivity of the mice after SNI surgery as measured by cold/hot plate test. One-way ANOVA with Dunnett’s multiple comparison tests. **(J)** Diagram of conditioned place preference/aversion test (left), the time that the mice spent in the chamber coupled with CNO treatment before (pre) and after (post) the conditioning (middle), and the CPP/A scores of the mice (right). RM two-way ANOVA with Tukey’s multiple comparison tests (middle) and one-way ANOVA with Dunnett’s multiple comparison tests (right). **(K)** Schematic summary showing the molecularly defined IT^Il1rapl2^, PT^Fezf2^ and CT^Foxp2^ neurons in pIC as discrete neuronal substrates with distinct connectivity, activities and functions in different dimensions of pain. \**P* < 0.05; \*\**P* < 0.01; ns, not significant.

The mice were first subjected to open field test (OFT) which revealed no significant differences in the distance traveled (**Fig. 6C**), indicating that inactivation of the IT^Il1rapl2^, PT^Fezf2^ or CT^Foxp2^ neurons does not significantly affect the locomotor activity of the mice. We then subjected the mice to von Frey test (VFT) to measure mechanical sensitivity. Compared to the control group, inactivation of the PT^Fezf2^ neurons (Fezf2-Gi) significantly decreased the response rate of the mice to the von Frey filament stimulation, whereas no significant change was observed in the Il1rapl2-Gi or Foxp2-Gi groups (**Fig. 6D**). These results indicate that inactivation of the PT^Fezf2^, but not the IT^Il1rapl2^ or CT^Foxp2^ neurons, significantly decreases mechanical sensitivity of the mice. To examine the roles of the neuron populations in regulating thermal pain sensitivity, the mice were subjected to cold plate test (CPT) and hot plate test (HPT) for cold and hot pain sensitivity, respectively (**Fig. 6E**). In both CPT and HPT, the Il1rapl2-Gi and Fezf2-Gi groups displayed significantly increased paw withdrawal latencies compared to the control group, while no significant difference was observed in the Foxp2-Gi group (**Fig. 6E**), indicating that inactivation of the IT^Il1rapl2^ or PT^Fezf2^, but not the CT^Foxp2^ neurons, significantly decreases cold and hot pain sensitivity. Together, these results demonstrate that the PT^Fezf2^ neurons modulate both mechanical and thermal pain sensitivity, and the IT^Il1rapl2^ neurons specifically modulate thermal pain sensitivity, whereas the CT^Foxp2^ neurons do not regulate pain sensitivity, suggesting the segregated processing pathways for mechanical and thermal pain in the pIC.

To investigate the roles of the neuron populations in chronic pain, SNI surgery was performed to induce chronic neuropathic pain in the mice, and the behavioral assays were performed four weeks later (**Fig. 6F**). The mice were subjected to OFT and again, no significant difference in traveled distance was observed (**Fig. 6G**), indicating inactivation of these neuron populations does not affect the locomotor activity of the mice with chronic neuropathic pain. We next examined the roles of the neuron populations in regulating mechanical allodynia by VFT and dynamic brush test (**Fig. 6H**). Compared to the control group, significant decrease in the response rate in VFT and dynamic score in brush test were observed in the Fezf2-Gi group, but not the Il1rapl2-Gi or Foxp2-Gi group (**Fig. 6H**), indicating that inactivation of the PT^Fezf2^, but not the IT^Il1rapl2^ or CT^Foxp2^ neurons, significantly relieves punctate and dynamic mechanical allodynia. The mice were also subjected to CPT and HPT to measure their cold and hot pain sensitivity, respectively (**Fig. 6I**). We found that the Il1rapl2-Gi and Fezf2-Gi groups, but not the Foxp2-Gi group, showed significantly increased paw withdrawal latencies in both the CPT and HPT (**Fig. 6I**), indicating that inactivation of the IT^Il1rapl2^ or PT^Fezf2^, but not the CT^Foxp2^ neurons, significantly decreases thermal pain sensitivity in mice with chronic neuropathic pain. To examine the roles of the neuron populations in regulating the negative affective valence of neuropathic pain, we subjected the mice to conditioned place preference/aversion (CPP/A) test (**Fig. 6J**), where the mice were treated with saline in one chamber and CNO in another chamber for chemogenetic inactivation. After three days of training, the Fezf2-Gi group spent significantly more time in the chamber paired with CNO treatment and surprisingly, the Foxp2-Gi group showed a significantly decreased CPP/A score, while no significant difference was observed in the Il1rapl2-Gi group (**Fig. 6J**). These results indicate that inactivation of the PT^Fezf2^ or the CT^Foxp2^ neurons respectively alleviates or exacerbates the negative affects of chronic neuropathic pain.

Collectively, these results demonstrate the distinct roles of the PT^Fezf2^, IT^Il1rapl2^ and CT^Foxp2^ neurons in pain processing, revealing the discrete neuron populations and pathways within the IC underlying different dimensions of pain. Inactivation of the PT^Fezf2^ neurons decreases both mechanical and thermal sensitivity and relieves the negative affective valence of chronic neuropathic pain, while inactivation of the IT^Il1rapl2^ neurons specifically regulates the thermal pain sensitivity. In contrast, inactivation of the CT^Foxp2^ neurons does not alter pain sensitivity but instead exacerbates the negative affective valence of chronic neuropathic pain.

## Discussion

In this study, we used MERFISH to resolve the spatial patterns of gene expression, cellular architecture and projection connectivity in the IC and adjacent regions. The major neuron classes exhibit distinct projection patterns, but transcriptomic type did not map one-to-one onto target specificity. Instead, the relationship between transcriptomic identity and connectivity varied across topographic positions. We further mapped the discrete input-output neural networks of the major neuron classes and identified the cell-type-specific involvement and functions of the pIC in pain processing (**Fig. 6K**). Together, our results establish a multimodal framework for understanding the molecular, cellular and circuit architecture of the IC at spatially resolved single-cell resolution and reveal the distinct roles for molecularly defined neuron types in pain.

Our analysis revealed the transcriptomically distinct cellular populations and their spatial distributions in the IC and adjacent regions. The neuronal composition of the IC diverges sharply from the ventrally adjacent Pir region but transitions continuously into neighboring neocortical regions. Interestingly, several transcriptomic types show regional enrichment or depletion, and some vary gradually across cortical subregions, indicating a graded cellular architecture within the IC and adjacent cortices. For example, the L2/3 IT 1 and L6 CT 2 neurons are relatively enriched in the agranular IC and depleted in the granular IC and SSC, whereas L2/3 IT 2 and L6 CT 1 neurons show the opposite distribution. Consistent with these cellular gradients, transcriptome-wide analyses identified genes and biological pathways selectively enriched or depleted in the IC relative to adjacent regions. These findings indicate that the IC contains a distinct molecular and cellular architecture that transitions continuously across neighboring cortical territories and may underlie regional differences in connectivity and function.

Transcriptomic identity is thought to encode developmental programs that shape neuronal morphology, connectivity and function(*5*). However, previous studies in the cortex and hypothalamus reported limited correspondence between transcriptomic identity and projection specificity(*9–11, 13*), despite clear relationships in retinal circuits(*37*). Our comprehensive survey shows that neuronal projection patterns depend on transcriptomic identity and spatial topography.

The major neuron classes exhibit segregated projection patterns: IT neurons target intratelencephalic regions such as the cortex, striatum and amygdala; PT neurons mainly innervate specific thalamic and brainstem nuclei; and CT neurons primarily project to the thalamus, with additional projections to brainstem nuclei. Consistent with observations in prefrontal and motor cortex(*9, 13*), the IC shows no simple one-to-one mapping between transcriptomic and projection types. Instead, each transcriptomic type projects to multiple targets, and each target receives projections from multiple transcriptomic types. Individual transcriptomic types within a major class share overlapping projection profiles, but with distinct spatially dependent patterns. For instance, a downstream target receives projections from multiple transcriptomic types mainly located at one subregion but not the same transcriptomic types in other subregions (for example, projections to acIC and CeA). The target may also receive projections from distinct transcriptomic types in different spatial locations (for example, projections to pcIC and NAc). Interestingly, the NTS mainly receives projections from the PT neurons regardless of the spatial location. For individual transcriptomic types, the projection targets shift depending on the spatial location of the neurons. Collectively, our study comprehensively deciphers how the projection pattern is governed by transcriptomic identity and spatial location. These principles suggest that a limited set of transcriptomic states can diversify circuit wiring through local topographic adaptation, thereby supporting specialized circuits for diverse brain functions.

By tracing the brain-wide projections and monosynaptic inputs, we established that the IT^Il1rapl2^, PT^Fezf2^ and CT^Foxp2^ neurons in pIC are integrated into discrete neural networks. The IT^Il1rapl2^ neurons target the intratelencephalic regions including the DP, NAc, CPu, BLA and CeA, and the CT^Foxp2^ neurons predominantly innervate the subregions of thalamus, while the PT^Fezf2^ neurons project to both thalamus and brainstem strongly. These three neuron populations receive comparable proportions of inputs from the cortices and pallidum, while the IT^Il1rapl2^ neurons receive more inputs from the thalamus and less from the amygdala, PBN and DRN. These connectivity networks demonstrate that the pIC does not strictly adhere to either a biased input-segregated output architecture or an integration-and-broadcast architecture(*38*). Instead, parallel neuron classes integrate biased upstream inputs and broadcast to segregated downstream targets, while the individual transcriptomic types within a class share the inputs and project broadly to multiple overlapping downstream targets, consistent with a hybrid model of both the biased input-segregated output architecture and integration-and-broadcast architecture.

Our results support the idea that the cerebral cortex comprises discrete IT, PT and CT modules, which have distinct transcriptomes, connectivity, activity and behavioral functions. While the pIC has been well documented for its roles in pain regulation(*21, 27*), its diverse neuron types and broad connectivity across the brain make the understanding of its specific cellular basis of multidimensional functions in pain difficult. We showed that the IT^Il1rapl2^ neurons are specifically activated by noxious thermal, but not mechanical, stimuli. Consistently, silencing these neurons elevates thermal pain thresholds in both naïve and chronic neuropathic pain states without affecting mechanical sensitivity. Given that the pIC contains primary representations of skin temperature(*34*), the IT^Il1rapl2^ neurons likely form a dedicated cellular channel for thermal nociception, potentially operating through downstream projections to the prefrontal cortex, striatum, or amygdala. Intriguingly, although the pIC-amygdala pathway has been described in affective processing(*26, 28*), inactivating the IT^Il1rapl2^ neurons failed to alter affective valence, suggesting that other IT populations, such as L2/3 IT neurons, may provide redundant or primary contributions to affective pain processing.

Previous studies have established that the pIC can modulate not only the descending facilitatory pathway via the raphe nuclei(*27*) but also the neuronal activity of the spinal cord through direct projections(*25*). Our results reveal that these projections arise from the PT^Fezf2^ neurons, which respond robustly to both mechanical and thermal noxious stimulation. We show that the PT^Fezf2^ neurons are the most affected population in chronic neuropathic pain, inactivation of these neurons also significantly alleviates mechanical allodynia, thermal hyperalgesia, and the negative affective valence of chronic neuropathic pain. This broad therapeutic effect is likely mediated by the projections to nociceptive hubs in the brainstem or spinal cord. In sharp contrast, the CT^Foxp2^ neurons are not activated, but instead exhibit a trend of decreased activity, in response to noxious mechanical or thermal stimulation. Silencing these neurons does not alter pain sensitivity but significantly exacerbates the negative affective valence of chronic neuropathic pain. The CT^Foxp2^ neurons predominantly project to the thalamus, which forms a reciprocal loop projecting back to the cortex(*39, 40*). The L6 CT projections typically act as modulatory feedback pathways capable of sustaining persistent thalamic activity(*40*). Inactivating the CT^Foxp2^ neurons likely disrupts this regulatory corticothalamic feedback, altering pIC dynamics to amplify affective pain. Together, our results demonstrate the IT^Il1rapl2^, PT^Fezf2^ and CT^Foxp2^ neurons in pIC as discrete modules with distinct transcriptome, connectivity and functions, revealing the segregated processing pathways of different dimensions of pain within the pIC.

Collectively, our study comprehensively deciphers the single-cell spatial organization of the IC at molecular, cellular and circuit levels, and demonstrates that the transcriptome and spatial location govern the neuronal projection patterns. We reveal the discrete neural basis in pIC with distinct transcriptome, connectivity and functions in regulating different dimensions of pain, providing a mechanistic framework for segregated cortical pain processing and targets for precision therapy of chronic pain.

## Materials and Methods

### Animals

All experiments were conducted in accordance with the National Institutes of Health Guide for Care and Use of Laboratory animals and approved by the Institutional Animal Care and Use Committee (IACUC) of Boston Children’s Hospital and Harvard Medical School. Wild-type C57BL/6 mice (Jax, #000664), B6;129S4-*Fezf2*^tm1.1(cre/ERT2)Zjh^/J (Jax, #036296), and B6.Cg-*Foxp2^tm1.1(cre)Rpa^*/J mice (Jax, #030541) were obtained from Jackson Laboratory. The *Il1rapl2*-Cre mouse line was described previously(*41*). Mice were group-housed with littermates (3-5 mice/cage) with food and water *ad libitum*. Housing environment was maintained under a 12-hr light/dark cycle (light on from 7:00 to 19:00) with ambient temperature (23-25°C) and humidity (55-62%) automatically controlled. For the present study, mice aged 2-4 months were used, and animals were randomly allocated to experimental groups. To minimize stress to the mice and avoid cross-experimental confounding effects, different behavioral assays were performed on separate days.

### Surgeries

Stereotaxic procedures were performed using a small-animal stereotaxic instrument (David Kopf Instruments, model 940). General anesthesia was induced and maintained using isoflurane (1.5 concentration in oxygen; 0.8 l min^−1^). Body temperature was sustained using a homeothermic feedback heating pad, and ophthalmic ointment was applied to prevent corneal desiccation throughout the procedure. A small craniotomy was made over the target brain region using a dental drill. Viral constructs or CTBs were delivered using a glass micropipette (tip diameter 10–20 μm) connected to a Nanoject III nanolitre injector (Nanoject III, Drummond Scientific, #3-200-207). For CTB tracing with integrated MERFISH, each mouse was injected with CTB-647 (Biotium, #00069) and CTB-740 (Biotium, #29127) into two target regions, respectively. At each target site, 120 nl of virus or 200 nl of CTB was injected at a rate of 1 nl s^−1^ unless otherwise specified. The micropipette was left in place for 15 min post-injection to permit solution diffusion before slow withdrawal. Following pipette removal, the wound was sutured, and the mice recovered on a warm blanket before being returned to their home cages. Postoperative care included daily subcutaneous injections of meloxicam (Covetrus, #049756) for 3 days, with health monitoring continued for an additional 2 days.

The coordinates of injection sites are based on previous literature and The Mouse Brain in Stereotaxic Coordinates (third edition) relative to the bregma (anterior-posterior, mediolateral, dorsoventral axis in mm): aIC (2.10, 2.25, -3.20), pIC (-0.82, 3.75, -4.00), mPFC (1.94, 0.45, - 2.50), SSp (0.02, 2.00, -1.50), NAc (1.10, 1.20, -4.50), CPu (0.62, 1.85, -4.50), BLA (-1.06, 2.85, -4.75), CeA (-1.34, 2.40, -4.50), MD (-1.22, 0.50, -3.20), HYT (-1.34, 0.75, -5.40), MRN (-3.80, 1.25, -3.50), PAG (-4.04, 0.45, -2.75), PBN (-5.34, 1.25, -3.50), NTS (-6.84, 1.00, -4.50).

For SNI surgery, mice were anesthetized with ketamine, the right hindlimb was shaved above the knee, and the incision site was disinfected using iodine and isopropanol. Blunt dissection of the overlying muscles exposed the three terminal branches of the sciatic nerve. The parallel tibial and common peroneal nerve branches were tightly ligated using two distinct sutures, and the nerve segment between the ligatures was transected and excised. The sural nerve branch was strictly avoided and left entirely undisturbed throughout the procedure. Overlying muscle layers were then released, and the skin incision was closed with sutures. For the sham-operated control group, identical surgical exposures were performed to reveal the sciatic nerve, but the branches were neither ligated nor transected prior to wound closure. The mice were allowed to recover on a warm blanket before being returned to their home cages. Mechanical allodynia was confirmed by VFT at specified post-surgical time points. For the mice prepared for MERFISH analysis, the brain tissues were collected 4 weeks after the surgery and the VFT was performed one day before the harvest. On the day of tissue harvesting, all acute handling was strictly minimized, and the mice were transferred directly from their home cages to the euthanasia chamber, after which brains were rapidly extracted and immediately frozen.

### Public scRNA-seq data analysis

The mouse whole cortex single-cell transcriptomes data were downloaded from Allen Brain Map(*30*) (https://portal.brain-map.org/atlases-and-data/rnaseq/mouse-whole-cortex-and-hippocampus-10x). The gene expression matrix and cell metadata were loaded into R with SingleCellExperiment(*42*) (v1.28.1) package. We only used cells from insula region for all downstream analysis, by selecting cells with region label annotation of ‘AI’. Then the data were converted to Seurat(*43*) (v5.3.0) object. We used the cell cluster label from the cell metadata as the cell type identity. The marker genes for each cell type were calculated with FindAllMarkers function of Seurat.

### MERFISH probes design and construction

The initial candidate genes for probes design were composed of the marker genes of each cell type (with a priority for neurons), ion channels, receptors, neuropeptide genes and activity-related genes. Genes with no expression in any cell types were filtered. Then genes with top 1% high expression in any cells (from the public scRNA-seq data) and genes with bottom 1% low expression in any cells were removed. We designed a codebook of 24 bits Hamming code with Hamming weight of 4 and Hamming distance of 4, using the python package MERFISH_probe_design (https://github.com/xingjiepan/MERFISH_probe_design). We included 50 unused barcodes as blank controls. Next, we designed the target probes for each gene (up to 96 target probes per gene) with the target length of 30 bp per probe, using the MERFISH_probe_design package and following the principles described previously(*44*). Specifically, we required the GC content of probes between 40% and 70%, the melting temperature between 50 and 65 °C, and the target specificity at least 0.99. Probes with high cis/trans-complementarity were filtered out. We required no overlap among the probes for the same gene. For genes with multiple isoforms, the common regions among all the isoforms were used for probes design. Genes with less than 40 target probes that could be designed meeting the above requirements were removed. This resulted in 434 genes with proper target probes (table S1). Then, each probe was added one or two readout complement sequences at both the 5’ end and 3’ end according to the codebook. Each bit of the 24 bits barcodes corresponded to a unique readout sequence. The readout sequences were from a previous study(*44*) and provided in table S2. For readout probes corresponding to the odd bits of the barcode, the Alexa Fluor 750 Dye were conjugated to their 5’ end by a disulfide bond. For readout probes corresponding to the even bits of the barcode, the Cy5 Dye were conjugated to their 5’ end by a disulfide bond. Finally, the flanking PCR amplification handle sequences were concatenated at 5’ end (CGGCTCGCAGCGTGTAAACG) and 3’ end (GCGTGGAGGGCATACAACG) for each probe. The final designed probe library for the 434 genes was provided in table S3.

The amplification and construction of the probe library was performed as described previously(*45*). Briefly, the synthesized probe library was amplified with limited-cycle PCR using a qPCR machine to monitor the amplification process and stop before reaching the plateau stage. Then the amplified DNA were purified with column and in vitro transcription was performed. The in vitro transcribed RNA was purified with beads and used as the template for reverse transcription. The products from the reverse transcription were processed through alkaline hydrolysis then purified via beads to obtain the final probe library.

### MERFISH imaging system

The imaging system for MERFISH was custom-built with the Nikon Ti2-E microscope, Lumencor CELESTA light engine and Hamamatsu ORCA-Fusion CMOS camera. The fluidics system with the Bioptechs FCS2 chamber was assembled as described in previously(*46*). The fluidics system was controlled automatically by the open-source software (https://github.com/ZhuangLab/storm-control). All the buffers used in the fluidics system were made according to the protocols described previously(*45*) and degassed for 10 min under vacuum prior to use. The software coordinating both the imaging system and the fluidics system to automatically perform MERFISH data acquisition was custom-developed. We used the Nikon CFI Plan Apochromat Lambda D 10X objective to define the imaging regions (field of view, FOV). The Nikon CFI Plan Apochromat Lambda D 60X Oil objective was used for MERFISH imaging. The multiple rounds of readout probe hybridization, imaging and fluorescence cleavage process were executed as described previously(*47*), for a total of 12 rounds for the 24-bit barcodes. For each FOV, we imaged 7 z-stack planes with a step of 1.5 μm. The 405 channel was used to image the DAPI and the 477 channel was used to image the total polyA RNA signals only in the first round of imaging, both of which were used for cell segmentation. The 545 channel was used to image the beads (Invitrogen, #F8800) coated onto the coverslip, which was used to align the images across different rounds of imaging. The 748 and 637 channel were used for the readouts imaging of the two bits in the barcodes per imaging round.

### MERFISH sample preparation

The samples for MERFISH imaging were prepared following the description in previous studies(*47–49*). The 40 mm diameter round coverslips (Bioptechs, #40-1313-0319) for mounting the samples were cleaned and silanized as described previously(*45*). Then the coverslips were coated with 0.1 mg/ml Poly-D-Lysine (Gibco, #A3890401) and 1:20,000 fluorescent beads (Invitrogen, #F8800) for 30 min, dried for 1 h, fixed with 4% PFA for 10 min, washed with pure water for 3 times and dried for 1 h under room temperature.

The mice were euthanized with CO_2_ and the brain tissues were quickly extracted, and immediately frozen in Optimal Cutting Temperature compound (OCT; Sakura, #4583) in a dry ice and ethanol slurry. The tissues were stored in -80 °C until sectioning. The fresh frozen samples were balanced in a cryostat (Leica, #3050S) prechilled to -18 °C for 1 hour, and then coronal slices were sectioned. Slices were collected for inspection under microscope and compared with the reference atlas to confirm the anatomic landmarks until the aIC (Bregma +2.0 mm) region was reached. A serial set of coronal slices (10 μm of thickness) were sectioned with the IC and adjacent areas dissected. Before collecting each slice onto the treated coverslip for imaging as described above, the previous one was inspected to confirm the alignment to the reference atlas. The slices were collected approximately every 200 μm of interval for one of the WT male mice (16 slices in total) and every 400 μm of interval for all other mice (8 slices for each mouse). Each coverslip accommodates 4 slices. For the sham and SNI mice, a pair of mice were sectioned at the same time, and the slices from both groups of mice were collected to the same coverslips.

After sectioning, the samples on the coverslips were thawed for 5 min at room temperature, fixed with 4% PFA for 10 min and washed twice with 1× PBS. The samples were stored in 70% ethanol at 4 °C for 24 h to permeabilize the cells. Next, the samples were washed twice with 2× SSC and once with encoding probe wash buffer (30% formamide in 2× SSC) for 5 min. Then the coverslip was inverted onto a 50 μl encoding probe buffer droplet on the flat parafilm in a petri dish. The encoding probe buffer contained a total of 3 μM encoding probes, 2 μM polyA-anchor probes, 30% formamide, 0.1% yeast tRNA, 10% dextran sulfate and 2× SSC. The coverslip was incubated at 37 °C for 36 h in a small humidified box in an incubator. Next, the coverslips were washed twice with the encoding probe wash buffer at 47 °C for 30 min. Then the samples were embedded in a thin polyacrylamide gel by inverting the coverslip onto a 65 μl gel solution on a gel-slick coated glass slides and incubated at room temperature for 2 h to allow the gel to polymerize. The gel embedded coverslip was carefully peeled up, put into the digestion buffer (2× SSC, 2% SDS, 0.25% Triton X-100, 1% proteinase K) and incubated at 37 °C for 48 h. After digestion, the coverslip was washed with 2× SSC at room temperature for 30 min on a rocking platform, and this wash was repeated four times. Finally, the samples were stored in 2× SSC at 4 °C before MERFISH imaging on the machine described above.

Immediately before MERFISH imaging, the samples on the coverslip were incubated in the hybridization buffer with the two readout probes of the first two bits for the first-round imaging (final concentration: 3nM each) and the readout probe (conjugated to Alexa Fluor 488 dye, 3nM) complementary to the readout sequence on the polyA-anchor, at room temperature for 15 min. Then the buffer was changed to the hybridization buffer with 0.4% 1 mg/ml DAPI solution, and incubated at room temperature for 10 min. Finally, the samples on the coverslip were washed with 2× SSC and used for imaging.

### MERFISH data analysis

The raw imaging data of MERFISH were decoded using the MERlin (https://github.com/emanuega/MERlin) pipeline with custom modifications to adapt to our custom-built machine as described above. We employed the Cellpose(*50*) software for cell segmentation based on the DAPI and total polyA RNA signals. The decoding results including the cell-by-gene count matrix, cell centroid coordinates and cell boundaries were loaded into the R package Seurat (v5.3.0) for downstream analysis. We only kept the cells with at least 60 RNAs and at least 20 different genes detected in the cell.

The raw counts were normalized using SCTransform. Principal components were calculated with the RunPCA function and UMAP embeddings were generated with the RunUMAP function. Single cells were clustered using FindNeighbors and FindClusters with a resolution of 4. The cell type of each cell was first automatically annotated with reference mapping using the FindTransferAnchors and TransferData functions, leveraging the public scRNA-seq data of insula cortex from the Allen Brain Map as the reference. For cells in a given cluster, the most frequent cell type annotated from the reference mapping was used as the cell type for that cluster. Then the cell type annotation for each cluster was manually checked and adjusted based on the marker genes of each cluster. The marker genes of each cluster were calculated with the FindAllMarkers function. The cells from different sample slices were integrated with the Harmony(*51*) method. To define the subtypes of each cell type, the cells in this cell type were taken out separately and reperformed the FindNeighbors and FindClusters with a resolution of 0.1. The average expression levels of genes in each cluster were calculated with the AverageExpression function. The spatial niche analysis was performed using the BuildNicheAssay function. To perform whole transcriptome imputation for each cell in the MERFISH data, we leveraged the reference mapping again with the public scRNA-seq data and imputed the expression levels for all the genes in each cell using the FindTransferAnchors and TransferData functions.

To register each sample slice to the Allen Brain Atlas CCF v3 coronal section(*52*), we used the python package abc_atlas_access (https://github.com/AllenInstitute/abc_atlas_access) to obtain the CCF v3 annotation. We implemented a python helper function to assist the manual adjustment to align each MERFISH slice to its closest matching CCF section based on the spatial border shapes and the spatial distribution of cell types. Then each cell was assigned with its anatomical structure after the registration. We used the anatomy structure of AId, AIp, AIv, CLA, GU and VISC as the broad insular cortex region.

The ARG score for the cells in control and SNI samples was calculated by averaging the z-score normalized expression level of the five genes: *Fos*, *Arc*, *Npas4*, *Nr4a3* and *Homer1*. The prioritization of cell types involved in pain condition was calculated with the Augur(*33*) package in R, and the AUC (Area Under the Receiver Operating Characteristic Curve) was used to rank the cell types. The clusterProfiler(*53*) (v4.6.2) package in R was employed to perform the Gene Ontology (GO) enrichment analysis.

### CTB imaging followed by MERFISH imaging

The brain tissues were harvested 7 days after the CTB injection and immediately frozen, followed by storage in -80 °C until cutting. The samples were sectioned into 10 μm slices with careful alignment to the reference atlas and mounted onto the treated coverslips, then fixed with 4% PFA for 10 min, washed with 1× PBS twice and put in 70% ethanol at 4 °C for 24 h. Next, the samples on the coverslip were washed twice with 2× SSC and mounted in the fluidics chamber for CTB imaging. The imaging regions were defined using the 10X objective under bright field. We switched to the 60X oil objective to image the beads (545 channel) on the coverslip and the CTB signals (748 / 637 channels). Then the coverslip was carefully removed from the fluidics chamber and processed the same as normal MERFISH sample preparation starting from the encoding probe hybridization as described above. To perform the MERFISH imaging after the sample preparation, we manually aligned the sample to the same position in the fluidics chamber as that during the CTB imaging. We selected a FOV in the center region and checked its beads position and adjusted the potential rotation, x/y offset to match the beads position of this FOV during CTB imaging. After MERFISH imaging, the CTB imaging data were combined with the MERFISH imaging data by aligning images based on the beads images to achieve exact registration, using the ORB (Oriented FAST and Rotated BRIEF) image matching (rigid transform, rotation/translation) from the OpenCV (cv2) python package. The CTB signals in each cell were calculated in the MERlin decoding pipeline with the SumSignal module.

### Single molecule fluorescence *in situ* hybridization (smFISH)

Mice were euthanized by inhalation of CO_2_ and transcardially perfused with PBS and 4% paraformaldehyde (PFA) before the brains were collected and fixed in 4% PFA at 4 °C overnight, followed by dehydration in 30% sucrose solution for 3 days. The brains were then frozen in OCT and coronal sections (14 μm) were cut using a cryostat (Leica, #CM3050S). The smFISH was performed using a RNAscope Fluorescent Multiplex Assay kit (Advanced Cell Diagnostics,) following the manufacturer’s instruction. Briefly, the brain slices were fixed with 4% PFA and washed with PBS, then dehydrated with ethanol and treated with hydrogen peroxide. The samples were then treated with the target retrieval reagents and protease for 10 minutes, after which the samples were hybridized with the probes in 40°C for 2 hours. Then the signals were amplified using the reagents included in the kit, followed by the labeling of relative channels with the TSA fluorophores. The samples were treated with the included blocking buffer before proceeding to DAPI staining and imaging. Probes (Advanced Cell Diagnostics) used were as followed: *Il1rapl2* (#552341-C2); *Fezf2* (#313301-C3); *Foxp2* (#428791).

### Neural tracing

To label the projections of the IT^Il1rapl2^, PT^Fezf2^ and CT^Foxp2^ neurons, pAAV1-hSyn-DIO-mGFP-Synaptophysin-mRuby (Addgene, #71760) was unilaterally injected into the pIC of *Il1rapl2*-Cre, *Fezf2*-CreER, or *Foxp2*-Cre mice. The brain tissues were collected 3 weeks later for imaging and examination. For monosynaptic retrograde rabies tracing, 150 nl mixture of AAV helpers (rAAV2/9-EF1a-DIO-mCherry-TVA and rAAV2/9-EF1a-DIO-RVG) (BrainVTA, #PT0023 and #PT0207) was unilaterally injected into the pIC of *Il1rapl2*-Cre, *Fezf2*-CreER, or *Foxp2*-Cre mice. Four weeks later, the same mice were injected with 250 nl of rabies virus (RV-ENVA-ΔG-EGFP) (BrainVTA, #R01001) to the same location. The brain tissues were collected 7 days later for imaging and examination. The mice were perfused as described above and the brains were harvested. Serial coronal slices (30 μm) were sectioned and mounted sequentially onto glass slides, which were then stored in -20 °C before DAPI staining. The tissues were imaged with a Zeiss LSM800 confocal microscope or Keyence BZ-X810 microscope, and the images were processed and analyzed with ImageJ for quantifications.

### Fiber photometry

The neuronal activities of the IT^Il1rapl2^, PT^Fezf2^ and CT^Foxp2^ neurons in pIC were measured using fiber photometry as previously described(*41*). Following the injection of AAV1-hSyn-FLEX-GCaMP7s (Addgene, #104491-AAV1) unilaterally into the pIC of *Il1rapl2*-Cre, *Fezf2*-CreER, or *Foxp2*-Cre mice, an optical cannula (diameter: 200 μm; N.A., 0.37; length, 4.0 mm; Inper Inc.) was implanted 100 μm above the virus injection sites. Following 3 weeks of recovery, the fluorescent signals were acquired with an RZ10X fiber photometry system equipped with LED drivers, LEDs and photosensors (Tucker-Davis Technologies). GCaMP7s fluorescence was excited using a 465 nm LED, while a 405 nm LED served as an isosbestic control to monitor motion and bleaching artifacts. Emitted light was collected through a Mini Cube (Doric Lenses) and spectrally separated into an autofluorescence reference band (420–450 nm) and the functional GCaMP7s signal band (500–550 nm). To minimize photobleaching, the LED power at the fiber tip was restricted to approximately 20 μW. Continuous recordings were capped at a maximum duration of 35 min per session. Mouse behaviors were recorded with CCD cameras (Super Circuits, Austin, TX). The GCaMP signal data stream was acquired with Synapse software (Tucker-Davis Technologies, version 44132) and were exported, filtered, and analyzed with Matlab code provided by TDT offline data analysis tools (https://www.tdt.com/docs/sdk/offline-data-analysis/offline-data-matlab/fiber-photometry-epoch-averaging-example/).

During the recording, mice were connected to an optical cable and acclimated in the testing chamber for 30 minutes prior to any behavioral stimulation. For mechanical stimulation, the mice were first recorded without stimulation for 2 min and then a pinprick or a light touch (0.04 g von Frey filament) was applied to the plantar of the contralateral hindlimb for 6 times, with a minimal interval of 2 min. For the thermal stimulation, the mice were similarly recorded, and cold (0 °C), warm (30 °C) or hot (55 °C) water was kept in a vacuum flask and for each trial, one drop of water was applied to the mice through a syringe. 6 trials were applied to each mouse in total, with a minimal interval of 2 min. For air puff stimulation, a 1-s air puff was applied onto the face of the mice, and 6 trials were performed to each mouse with a minimal interval of 2 min.

The GCaMP data streams were exported and analyzed with MATLAB codes provided by TDT offline data analysis tools (https://www.tdt.com/docs/sdk/offline-data-analysis/offline-data-matlab/fiber-photometry-epoch-averaging-example/). The data streams were segmented into individual trials based on different events with a time window spanning 5 s pre-stimulus to 10 s post-stimulus. Photobleaching and movement artifacts were corrected by applying a polynomial linear fitting was applied to the isosbestic signals to align it to the GCaMP signals. All the trials were shown for the heatmaps and averaged to generate the averaged traces, which were presented using the mean and standard deviation. To quantify the GCaMP signals, the signals 5 s before and 5 s after stimulation were averaged for each mouse, and each data point represents one individual mouse recorded.

### Chemogenetic manipulation

To inactivate the activities of the IT^Il1rapl2^, PT^Fezf2^ and CT^Foxp2^ neurons in pIC, AAV5-hSyn-DIO-mCherry (Addgene, #50459) or AAV5-hSyn-DIO-hM4D(Gi)-mCherry (Addgene, #44362) was bilaterally injected into the pIC of *Il1rapl2*-Cre, *Fezf2*-CreER, or *Foxp2*-Cre mice. The behavioral assays were performed 3 weeks later, and the mice were intraperitoneally injected with CNO (2 mg/kg, Cayman, #16882) in saline 20 min before each behavioral test. The virus expression was confirmed by *post hoc* histological examination and the mice with off-target virus expression were excluded.

### Tamoxifen induction

To drive Cre-dependent recombination in *Fezf2*-CreER mice, animals were administered tamoxifen (Sigma-Aldrich, #T5648). Tamoxifen solutions were freshly prepared by dissolving the reagent in corn oil to a final concentration of 20 mg/ml by constant shaking for 24 h at room temperature within a container protected from light. Mice received intraperitoneal injections of tamoxifen (100 mg/kg) once every 2 days for a total of 3 doses, commencing the day following stereotaxic surgery. For the chemogenetic experiment, the *Fezf2*-CreER mice received tamoxifen injections, whereas the *Il1rapl2*-Cre and *Foxp2*-Cre mice were injected with corn oil without tamoxifen. Animal health was carefully monitored across the entire injection timeline.

### Behavioral assays

#### Open field test

Open field test was performed to measure the locomotor activity of the mice as previously described(*54*). Briefly, an eight-chamber (W x L x H: 27 cm x 27 cm x 24 cm, ENV-510 boxes) activity monitor system (Med Associates) was used for the open field test. After 20 min of acclimation, individual mice were placed in the center of the box and allowed to freely explore the chamber for 30 min. Locomotor activities of the mice were recorded and analyzed.

#### Von Frey test

A series of von Frey filaments (ranging from 0.04 – 2.0 g, Stoelting) were used to measure the mechanical sensory threshold or response rates as previously described(*29, 55*). Briefly, the mice were placed in a transparent plastic chamber on an elevated wire mesh floor and allowed to move freely. After 20 min of acclimation, the von Frey filaments were applied to the mid-plantar surface of the hindpaw through the mesh floor and kept for 2 s, or until a positive response (paw withdrawal, flinching, licking or shaking) was elicited. To measure the response rates, the von Frey filaments were applied in the order of ascending forces (5 times for each filament, 30 s interval). To avoid potential false-positive responses associated with spontaneous movement, the measurement was conducted again if a positive response was immediately followed by other voluntary behaviors such as locomotion, exploratory or grooming behaviors.

#### Cold/Hot plate test

The cold/hot plate test was used to measure thermal nociception of the mice. After being acclimated in the testing room for 20 min, the mice were placed on a plate (Ugo Basile) set at 0 °C (cold) or 52 °C (hot) and the latencies to flinching or licking of hindpaws were measured. All animals were tested 3 times with a minimum interval of 30 min. To avoid tissue injury, a cutoff latency was set at 45 s (cold) or 30 s (hot) for the test.

#### Conditioned place preference/aversion

The conditioned place preference test was used to measure the affective valences of pain or pain relief as previously described(*55*). The testing apparatus (Med Associates) consisted of two main chambers featuring distinct contextual environments (rod-style versus grid-style flooring) and was equipped with an array of infrared photobeam detectors to continuously track animal positioning and locomotor trajectories. On day 1, individual mice were placed in the apparatus and allowed to freely explore both chambers for 30 min (pre-test). On days 2–4, mice underwent two separate 30-min training sessions daily, maintaining a minimum inter-session interval of 6 h. During the morning sessions, animals were administered saline and confined to the chamber with grid floor (unconditioned chamber) for 30 min. During the afternoon sessions, mice received CNO treatment and were placed in the opposite chamber with rod floor (conditioned chamber). On day 5 (post-test), the mice were again permitted to freely explore the entire apparatus for 30 min. To calculate the CPP score, the time that mice spent in the conditioned compartment during pre-test was subtracted from the one during post-test.

### Statistics

Mice were randomly allocated to experimental groups, and the investigators were blind to the group identities during data collection or outcome assessments. Sample sizes were not predetermined using statistical power calculations, but our sample sizes are similar to those reported in previous publications. Mice that, after histological inspection, had the location of the virus expression or optic fiber implantation outside the area of interest were excluded. Statistical analyses were performed using GraphPad Prism (version 10) unless otherwise specified. Nonparametric tests, student’s t-test, one-way or two-way ANOVA tests with *post hoc* multiple comparisons were applied to determine statistical differences. All data are presented as means ± SEM. Statistical significance was set at \**P* < 0.05, \*\**P* < 0.01, \*\*\**P* < 0.001.

## Acknowledgments

We thank the support of Mouse Behavior Core of Harvard Medical School and its director Dr. Barbara Caldarone. We thank Dr. Chao Zhang for help in public scRNA-seq analysis.

## Funding

National Institutes of Health grant 1R01DA062669 (YZ)

Howard Hughes Medical Institute (YZ)

Open Philanthropy Foundation (YZ)

## Author contributions

Y.Z. conceived and supervised the project; G.X. and M.W. designed the experiments; M.W. built the MERFISH equipment and developed the automation software; G.X. and M.W. performed the MERFISH experiments; M.W. performed bioinformatic analysis; G.X. and Y.L. performed the biological experiments; All authors interpreted the data; G.X., M.W. and Y.Z. wrote the manuscript; All authors reviewed the manuscript.

## Data, code, and materials availability

All the MERFISH data generated in this study have been deposited to the NCBI Gene Expression Omnibus (GEO) with accession number GSE329718. The processed data in R Seurat object can be downloaded at https://doi.org/10.6084/m9.figshare.32193567. The code used in this study is available at GitHub: https://github.com/YiZhang-lab/Insula_MERFISH.

