## Supplementary materials for "Spatially resolved cellular and circuit architecture of the insular cortex controlling different dimensions of pain"

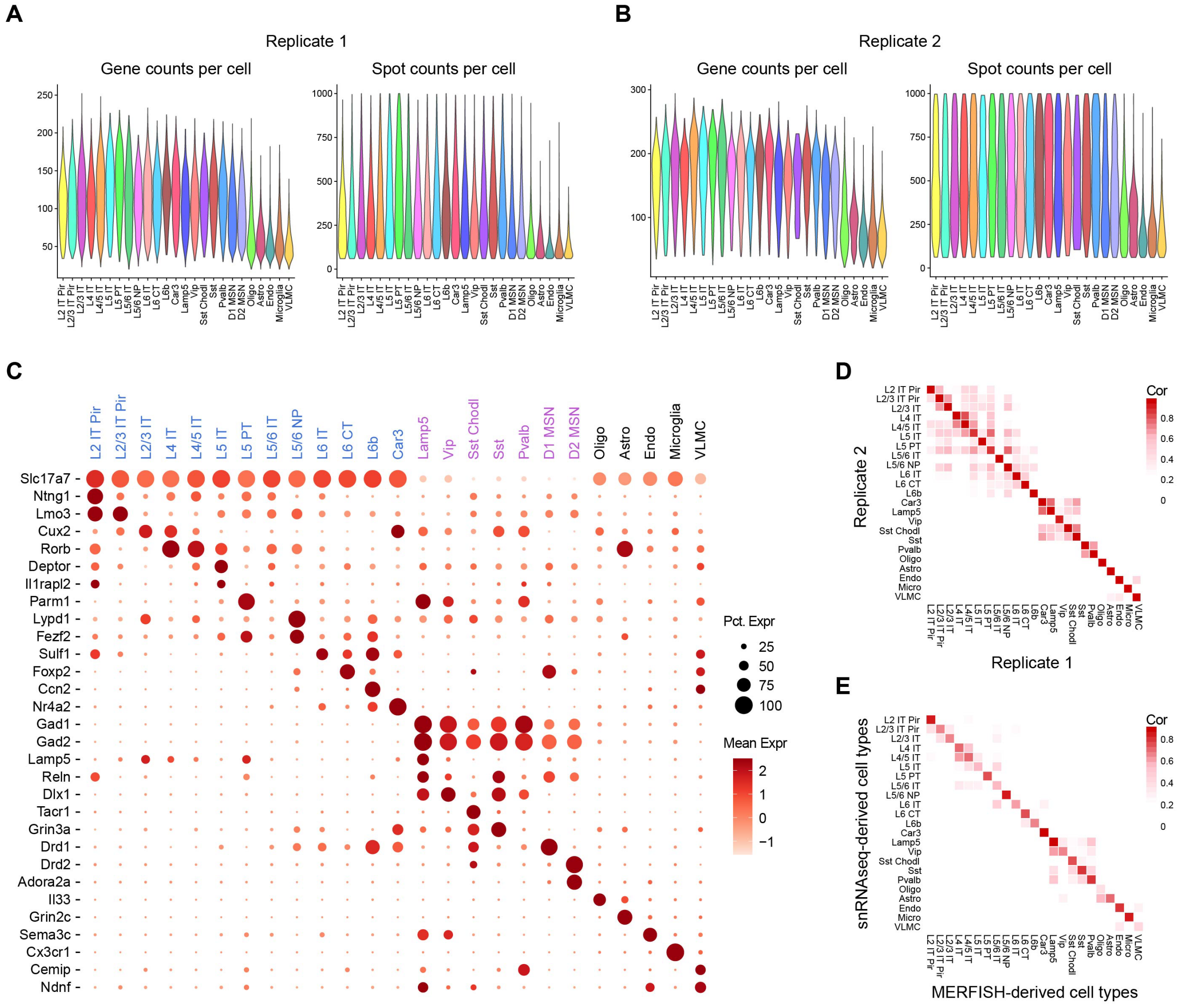

**Fig. S1. Quality control of MERFISH data. (A, B)** Violin plot showing the distribution of gene detection and mRNA molecule detection for the two biological replicates of two male mice. **(C)** Expression of marker genes for the transcriptomic cell types identified by MERFISH. The excitatory neurons, inhibitory neurons and non-neurons are colored in blue, purple or black, respectively. **(D)** Heatmap showing the gene expression correlation of the cell types between the two biological replicates. **(E)** Heatmap showing the gene expression correlation of the cell types from MERFISH and public scRNA-seq data.

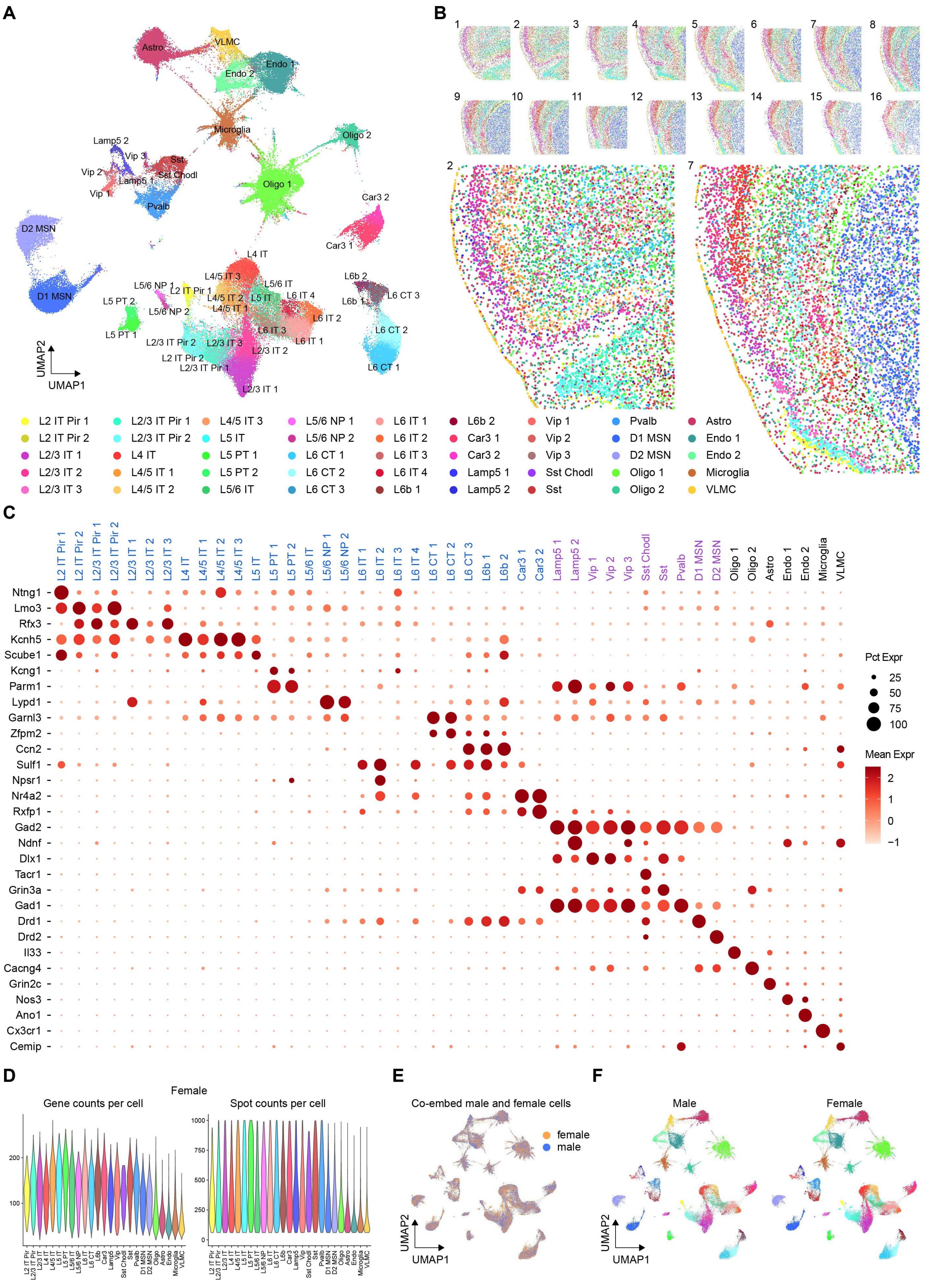
A
Astro
VLMC
Endo 1
Endo 2
Microglia
Oligo 2
Oligo 1
Lamp5 2
Vip 3
Sst
Sst Chodl
Vip 2
Lamp5 1
Vip 1
Pvalb
Car3 2
Car3 1
D2 MSN
D1 MSN
L4 IT
L4/5 IT 3
L5/6 IT
L5/6 NP 1
L2 IT Pir 1
L6 IT 4
L4/5 IT 2
L5 IT
L4/5 IT 1
L5/6 NP 2
L5 PT 2
L6 IT 2
L6 IT 3
L6 IT 1
L5 PT 1
L2/3 IT Pir 2
L2/3 IT 3
L2/3 IT 2
L2 IT Pir 2
L2/3 IT Pir 1
L2/3 IT 1
L6b 2
L6b 1
L6 CT 3
L6 CT 2
L6 CT 1
UMAP2
UMAP1
B
2
7
L2 IT Pir 1
L2 IT Pir 2
L2/3 IT 1
L2/3 IT 2
L2/3 IT 3
L2/3 IT Pir 1
L2/3 IT Pir 2
L4 IT
L4/5 IT 1
L4/5 IT 2
L4/5 IT 3
L5 IT
L5 PT 1
L5 PT 2
L5/6 IT
L5/6 NP 1
L5/6 NP 2
L6 CT 1
L6 CT 2
L6 CT 3
L6 IT 1
L6 IT 2
L6 IT 3
L6 IT 4
L6b 1
L6b 2
Car3 1
Car3 2
Lamp5 1
Lamp5 2
Vip 1
Vip 2
Vip 3
Sst Chodl
Sst
Pvalb
D1 MSN
D2 MSN
Oligo 1
Oligo 2
Astro
Endo 1
Endo 2
Microglia
VLMC
C
L2 IT Pir 1
L2 IT Pir 2
L2/3 IT Pir 1
L2/3 IT Pir 2
L2/3 IT 1
L2/3 IT 2
L2/3 IT 3
L4 IT
L4/5 IT 1
L4/5 IT 2
L4/5 IT 3
L5 IT
L5 PT 1
L5 PT 2
L5/6 IT
L5/6 NP 1
L5/6 NP 2
L6 IT 1
L6 IT 2
L6 IT 3
L6 IT 4
L6 CT 1
L6 CT 2
L6 CT 3
L6b 1
L6b 2
Car3 1
Car3 2
Lamp5 1
Lamp5 2
Vip 1
Vip 2
Vip 3
Sst Chodl
Sst
Pvalb
D1 MSN
D2 MSN
Oligo 1
Oligo 2
Astro
Endo 1
Endo 2
Microglia
VLMC
Ntng1
Lmo3
Rfx3
Kcnh5
Scube1
Kcng1
Parm1
Lypd1
Garnl3
Zfpm2
Ccn2
Sulf1
Npsr1
Nr4a2
Rxfp1
Gad2
Ndnf
Dlx1
Tacr1
Grin3a
Gad1
Drd1
Drd2
Il33
Cacng4
Grin2c
Nos3
Ano1
Cx3cr1
Cemip
Pct Expr
25
50
75
100
Mean Expr
2
1
0
−1
D
Female
Gene counts per cell
Spot counts per cell
E
Co-embed male and female cells
female
male
UMAP2
UMAP1
F
Male
Female
UMAP2
UMAP1

**Fig. S2. The spatial organization of the cellular subtypes in the IC and adjacent regions. (A)** UMAP visualization of the cellular subtypes revealed by MERFISH, integrating all the cells from the two biological replicates. The cells are color-coded by their subtype identities. **(B)** An overview of all spatial maps of the cellular subtypes in all coronal slices (top) from one male mouse, and the enlarged spatial maps for the anterior or posterior IC and adjacent regions (bottom). The cells are color-coded by their subtype identities. **(C)** Expression of marker genes for the cellular subtypes. The excitatory, inhibitory and non-neuron subtypes are colored in blue, purple or black, respectively. **(D)** Violin plot showing the distribution of the gene detection (left) and mRNA molecule detection (right) in the brain sections from a female mouse. **(E)** UMAP showing the co-embedding of the male and female dataset. **(F)** UMAP visualization of the cellular subtypes of the male (left) or female (right) mice. The cells are color-coded by their subtype identities.

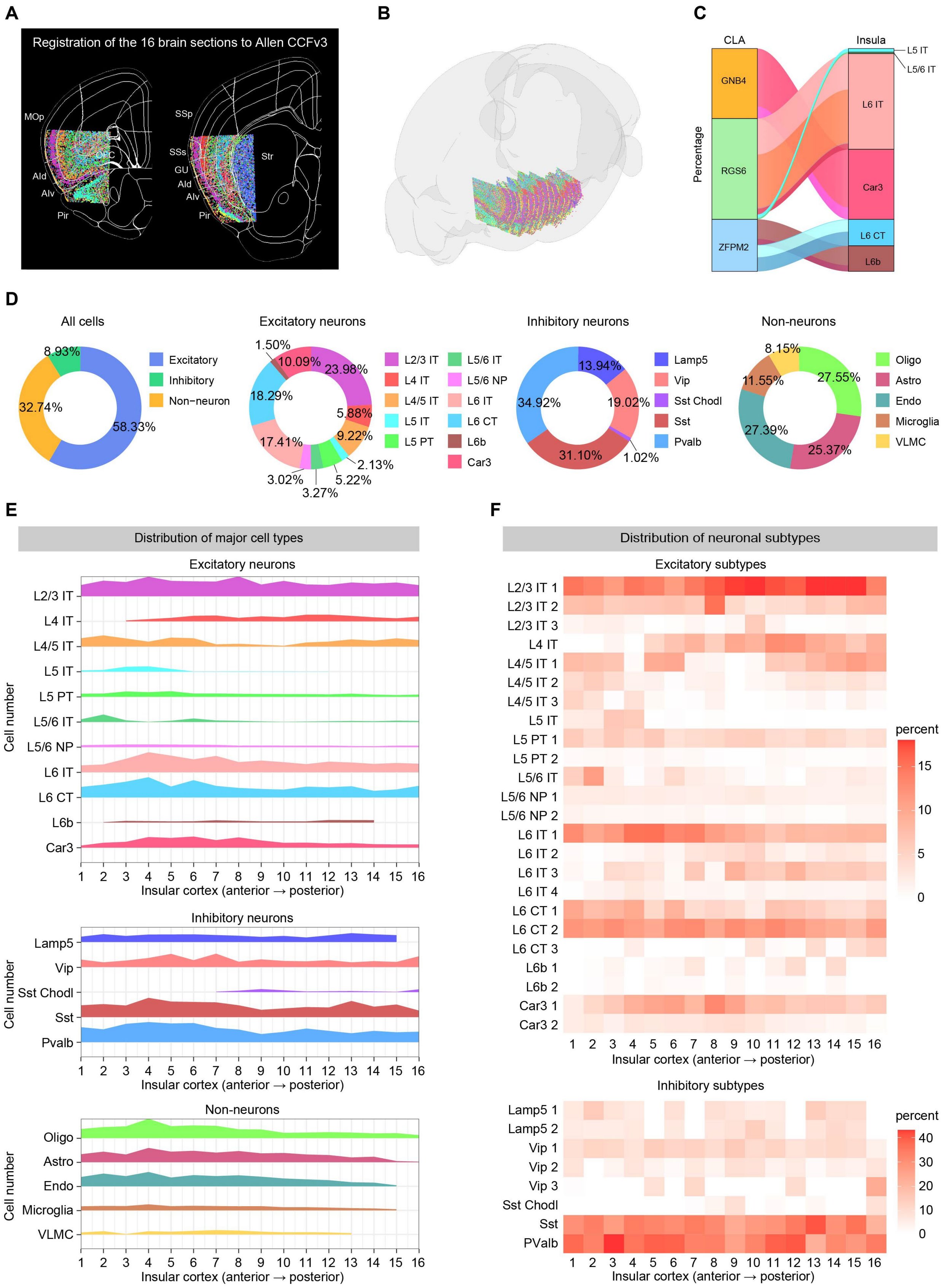
A
Registration of the 16 brain sections to Allen CCFv3
MOp
AId
AIv
Pir
SSp
SSs
GU
AId
AIv
Pir
Str
B
C
CLA
Insula
GNB4
RGS6
ZFPM2
L5 IT
L5/6 IT
L6 IT
Car3
L6 CT
L6b
Percentage
D
All cells
Excitatory
Inhibitory
Non−neuron
8.93%
32.74%
58.33%
Excitatory neurons
1.50%
10.09%
23.98%
18.29%
5.88%
9.22%
17.41%
2.13%
3.02%
5.22%
3.27%
L2/3 IT
L4 IT
L4/5 IT
L5 IT
L5 PT
L5/6 IT
L5/6 NP
L6 IT
L6 CT
L6b
Car3
Inhibitory neurons
13.94%
19.02%
34.92%
31.10%
1.02%
Lamp5
Vip
Sst Chodl
Sst
Pvalb
Non-neurons
8.15%
11.55%
27.55%
27.39%
25.37%
Oligo
Astro
Endo
Microglia
VLMC
E
Distribution of major cell types
Excitatory neurons
Inhibitory neurons
Non-neurons
Cell number
Insular cortex (anterior → posterior)
F
Distribution of neuronal subtypes
Excitatory subtypes
L2/3 IT 1
L2/3 IT 2
L2/3 IT 3
L4 IT
L4/5 IT 1
L4/5 IT 2
L4/5 IT 3
L5 IT
L5 PT 1
L5 PT 2
L5/6 IT
L5/6 NP 1
L5/6 NP 2
L6 IT 1
L6 IT 2
L6 IT 3
L6 IT 4
L6 CT 1
L6 CT 2
L6 CT 3
L6b 1
L6b 2
Car3 1
Car3 2
percent
Insular cortex (anterior → posterior)
Inhibitory subtypes
Lamp5 1
Lamp5 2
Vip 1
Vip 2
Vip 3
Sst Chodl
Sst
PValb
percent
Insular cortex (anterior → posterior)

**Fig. S3. Cellular composition and anterior-posterior distribution of the cell types in IC. (A)** Example MERFISH-derived cell-type spatial maps registered to the Allen CCFv3 for anatomical annotation. The cells are color-coded by their identities. **(B)** 3D reconstruction of the spatial maps of the entire imaged region after anatomical annotations for one male mouse. **(C)** Alluvial diagram showing the correspondence of the glutamatergic neuron types in claustrum and the IC. **(D)** Pie charts showing the proportions of the major cell types, excitatory neurons, inhibitory neurons and non-neurons in the IC. **(E)** Anterior-posterior distributions of the excitatory neurons (top), inhibitory neurons (middle) and non-neurons (bottom) in the IC. The Y axis indicates the cell numbers. **(F)** Heatmaps showing the anterior-posterior distributions of the subtypes of excitatory neurons (top) and inhibitory neurons (bottom) in the IC. The color bar indicates the proportions of the cell types in the region.

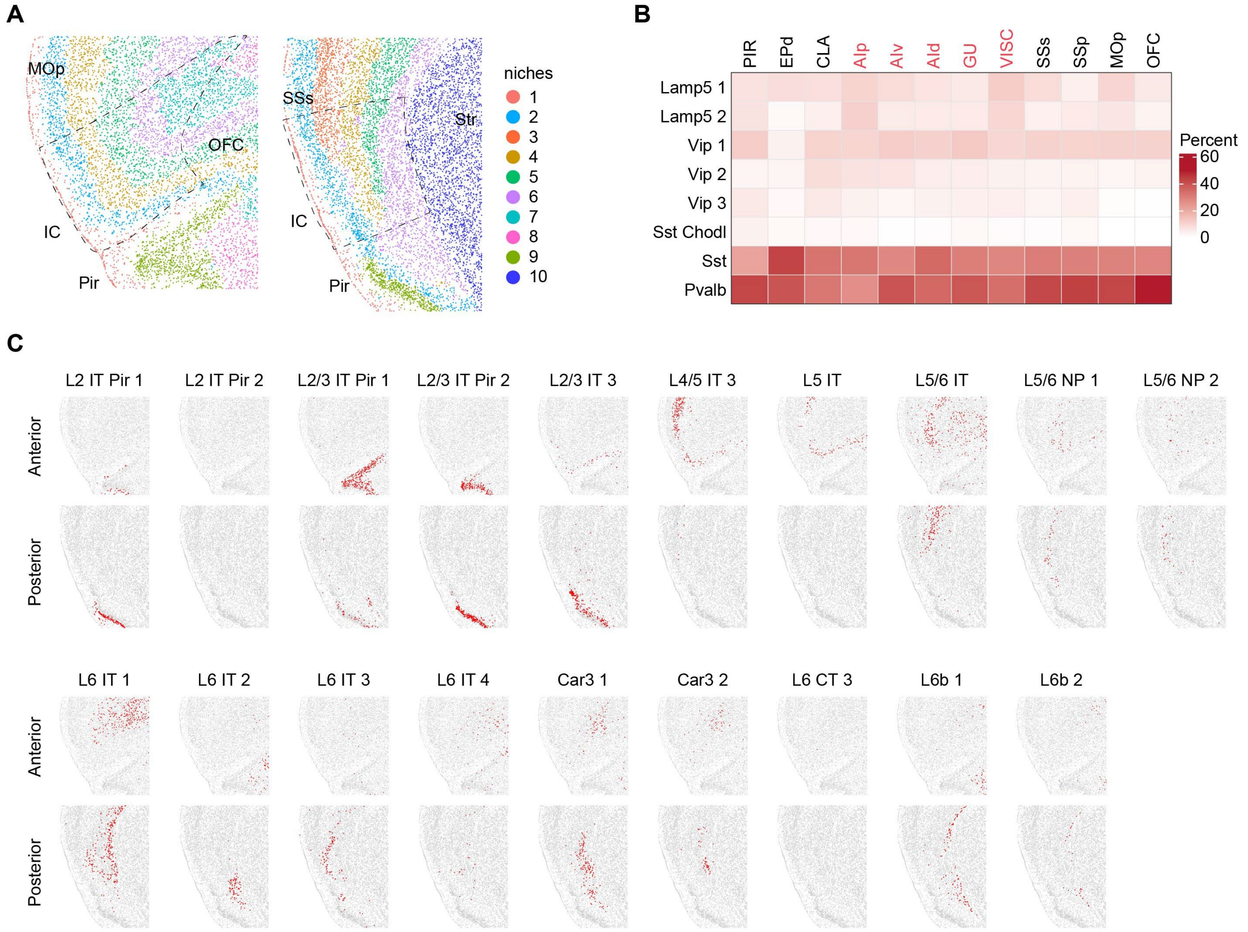

**Fig. S4. Spatial organization of the neuron subtypes in the IC. (A)** Spatial niches identified from the MERFISH data in the anterior (left) and posterior (right) IC. **(B)** Heatmap showing the proportion of inhibitory neuron subtypes in different anatomical regions. The subregions of IC are highlighted in red. **(C)** Spatial location of the excitatory neuron subtypes on the representative coronal slices for anterior and posterior parts of IC. Red dots represent the indicated subtypes.

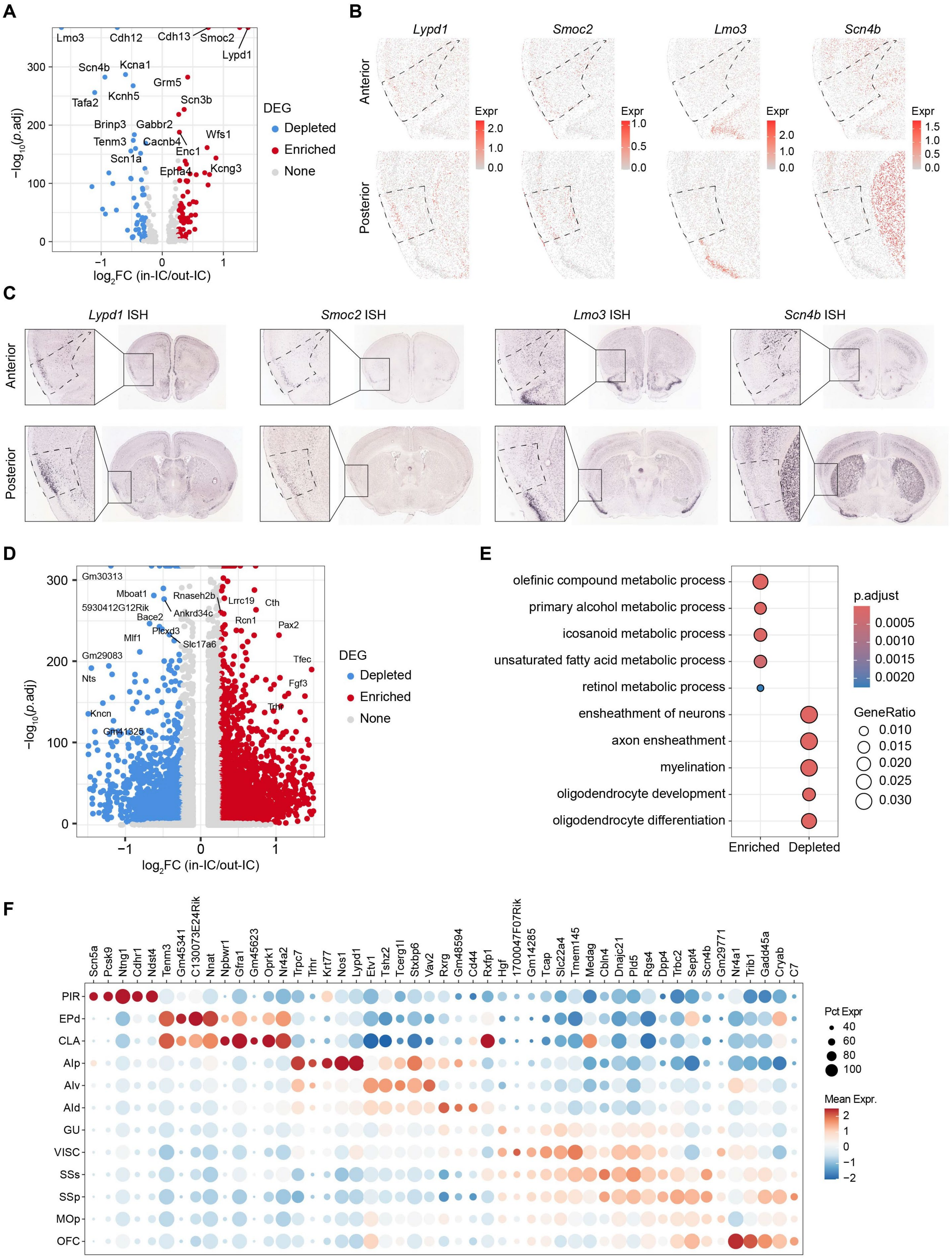

A
Lmo3
Cdh12
Cdh13
Smoc2
Lypd1
Scn4b
Kcna1
Grm5
Kcnh5
Tafa2
Scn3b
Brinp3
Gabbr2
Tenm3
Cacnb4
Wfs1
Enc1
Scn1a
Epha4
Kcng3
−log10(p.adj)
log2FC (in-IC/out-IC)
DEG
Depleted
Enriched
None
B
Lypd1
Smoc2
Lmo3
Scn4b
Anterior
Posterior
Expr
C
Lypd1 ISH
Smoc2 ISH
Lmo3 ISH
Scn4b ISH
Anterior
Posterior
D
Gm30313
Mboat1
5930412G12Rik
Rnaseh2b
Ankrd34c
Lrrc19
Cth
Bace2
Rcn1
Pax2
Plcxd3
Mlf1
Slc17a6
Gm29083
Tfec
Nts
Fgf3
Trh
Kncn
Gm41326
−log10(p.adj)
log2FC (in-IC/out-IC)
DEG
Depleted
Enriched
None
E
olefinic compound metabolic process
primary alcohol metabolic process
icosanoid metabolic process
unsaturated fatty acid metabolic process
retinol metabolic process
ensheathment of neurons
axon ensheathment
myelination
oligodendrocyte development
oligodendrocyte differentiation
Enriched
Depleted
p.adjust
0.0005
0.0010
0.0015
0.0020
GeneRatio
0.010
0.015
0.020
0.025
0.030
F
Scn5a Pcsk9 Ntng1 Cdhr1 Ndst4 Tenm3 Gm45341 C130073E24Rik Nnat Npbwr1 Gfra1 Gm45623 Oprk1 Nr4a2 Trpc7 Trhr Krt77 Nos1 Lypd1 Etv1 Tshz2 Tcerg1l Stxbp6 Vav2 Rxrg Gm48594 Cd44 Rxfp1 Hgf 1700047F07Rik Gm14285 Tcap Slc22a4 Tmem145 Medag Cbln4 Dnajc21 Pld5 Rgs4 Dpp4 Trbc2 Sept4 Scn4b Gm29771 Nr4a1 Trib1 Gadd45a Cryab C7
PIR
EPd
CLA
AIp
AIv
AId
GU
VISC
SSs
SSp
MOp
OFC
Pct Expr
40
60
80
100
Mean Expr.
2
1
0
−1
−2

**Fig. S5. Spatial organization of the gene expression in the IC. (A)** Volcano plot showing the DEGs that are enriched or depleted in the IC neurons relative to the neurons in the adjacent regions. Expression of genes enriched or depleted in IC are colored in red or blue, respectively. Threshold: adjusted p-value $< 0.01$ and fold change $\geq 1.2$. **(B)** Spatial expression of the representative enriched or depleted genes *Lypd1*, *Smoc2*, *Lmo3* and *Scn4b* in the representative anterior (top) and posterior (bottom) IC and adjacent regions derived from MERFISH data. **(C)** ISH data from Allen Brain Atlas showing the spatial expression of *Lypd1*, *Smoc2*, *Lmo3* and *Scn4b* in the representative anterior (top) and posterior (bottom) IC and adjacent regions with magnified images of the indicated regions. **(D)** Volcano plot showing the genes enriched or depleted in the IC after whole transcriptome imputing by correlating the MERFISH data and public scRNA-seq data. The enriched or depleted genes are colored in red or blue, respectively. Threshold: adjusted p-value $<$ 0.01 and fold change $\geq 1.2$. **(E)** Gene Ontology (GO) enriched terms for the IC enriched and depleted genes identified in panel **(D)**. **(F)** Heatmap showing the marker genes of different anatomical regions. The marker genes were identified from imputed whole transcriptome.

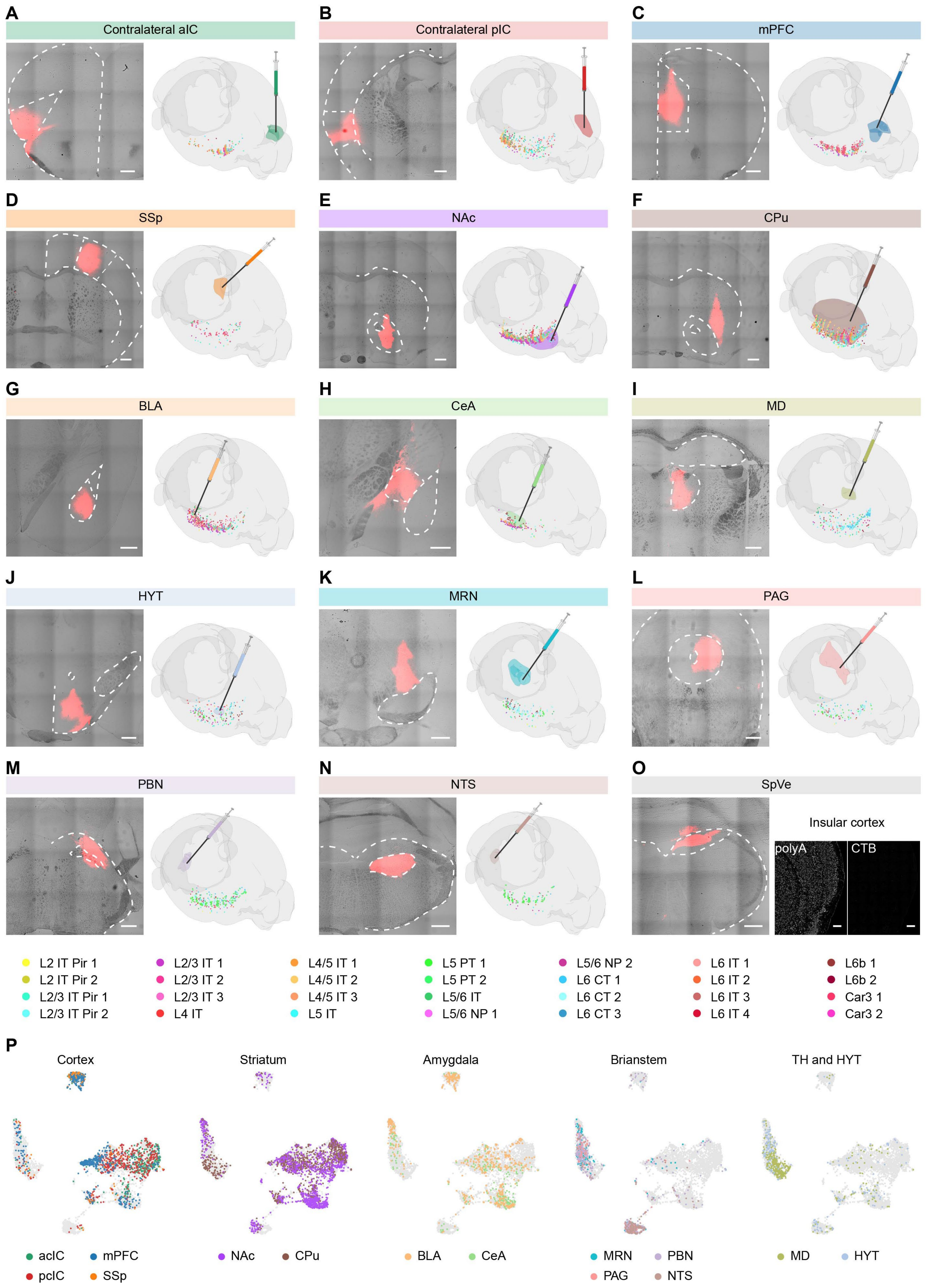

A
Contralateral aIC
B
Contralateral pIC
C
mPFC
D
SSp
E
NAc
F
CPu
G
BLA
H
CeA
I
MD
J
HYT
K
MRN
L
PAG
M
PBN
N
NTS
O
SpVe
Insular cortex
polyA
CTB
L2 IT Pir 1
L2 IT Pir 2
L2/3 IT Pir 1
L2/3 IT Pir 2
L2/3 IT 1
L2/3 IT 2
L2/3 IT 3
L4 IT
L4/5 IT 1
L4/5 IT 2
L4/5 IT 3
L5 IT
L5 PT 1
L5 PT 2
L5/6 IT
L5/6 NP 1
L5/6 NP 2
L6 CT 1
L6 CT 2
L6 CT 3
L6 IT 1
L6 IT 2
L6 IT 3
L6 IT 4
L6b 1
L6b 2
Car3 1
Car3 2
P
Cortex
Striatum
Amygdala
Brianstem
TH and HYT
acIC
mPFC
pcIC
SSp
NAc
CPu
BLA
CeA
MRN
PBN
PAG
NTS
MD
HYT

**Fig. S6. Integrated MERFISH and retrograde neural tracing for the IC. (A-N)** Injection sites of the CTB (red) for the 14 indicated projection targets (left), and the diagram showing the injection site in the brain as well as the reconstructed 3D model of the labeled neurons in the IC and adjacent regions (right). The cells are color-coded by their identities as indicated at the bottom. Scale bars, 200 μm. **(O)** Injection of CTB into the SpVe (left), all polyA mRNA signals of the cells (middle) and no detectable signal of CTB in the IC (right). Scale bars, 200 μm. **(P)** UMAP visualization of the IC neurons retrogradely labeled from the different projection targets. The cells are color-coded by the projection targets.

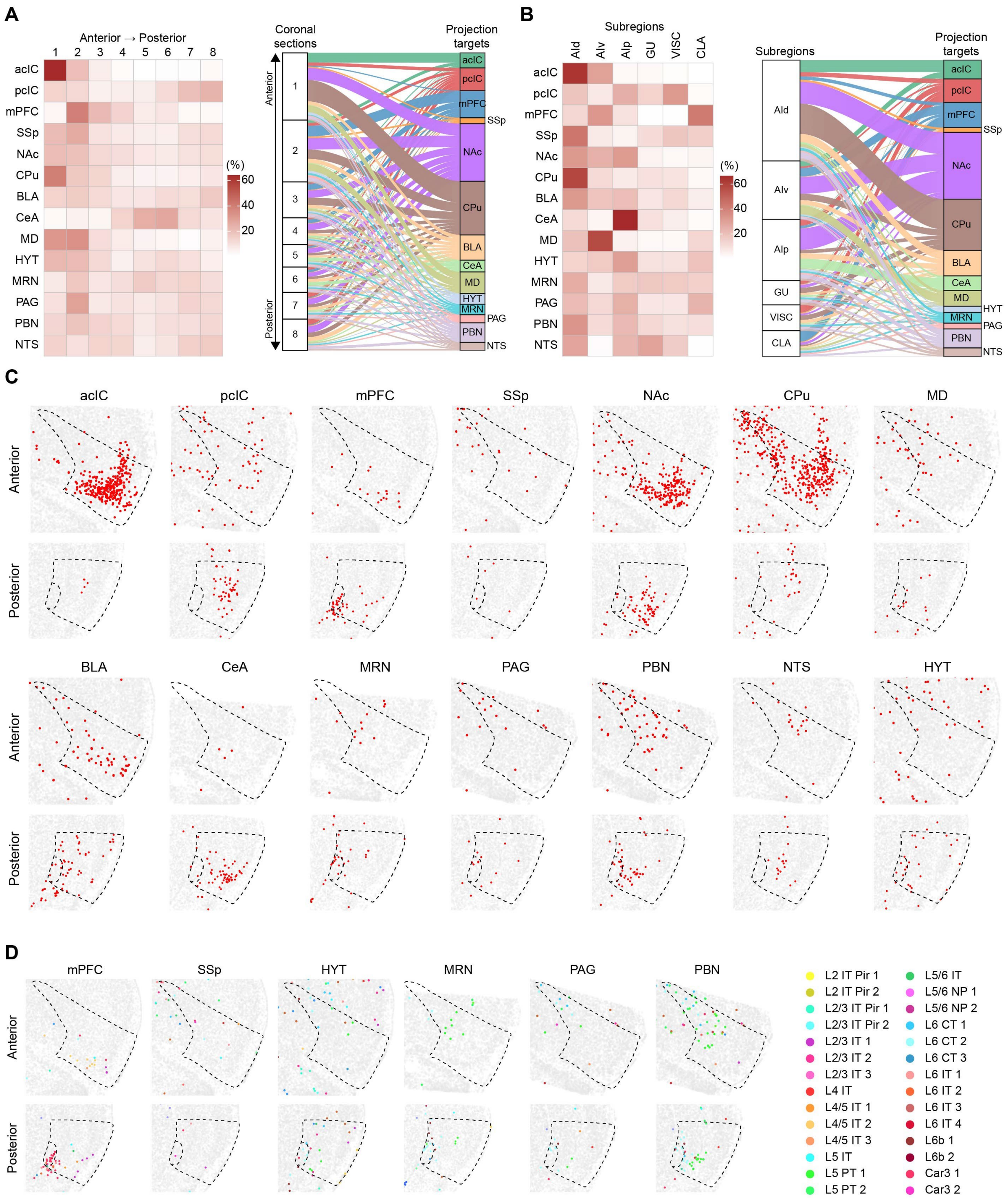
A
Anterior → Posterior
1 2 3 4 5 6 7 8
acIC
pcIC
mPFC
SSp
NAc
CPu
BLA
CeA
MD
HYT
MRN
PAG
PBN
NTS
(%)
60
40
20
Coronal sections
Projection targets
Anterior
Posterior
B
Subregions
AId AIv AIp GU VISC CLA
Projection targets
C
acIC pcIC mPFC SSp NAc CPu MD
BLA CeA MRN PAG PBN NTS HYT
D
mPFC SSp HYT MRN PAG PBN
L2 IT Pir 1
L2 IT Pir 2
L2/3 IT Pir 1
L2/3 IT Pir 2
L2/3 IT 1
L2/3 IT 2
L2/3 IT 3
L4 IT
L4/5 IT 1
L4/5 IT 2
L4/5 IT 3
L5 IT
L5 PT 1
L5 PT 2
L5/6 IT
L5/6 NP 1
L5/6 NP 2
L6 CT 1
L6 CT 2
L6 CT 3
L6 IT 1
L6 IT 2
L6 IT 3
L6 IT 4
L6b 1
L6b 2
Car3 1
Car3 2

**Fig. S7. Spatial organization of the retrogradely labeled neurons in the IC and adjacent regions. (A)** Heatmap (left) and Sankey plot (right) showing the anterior-posterior distribution of the retrogradely labeled neurons in the IC and adjacent regions from the different projection targets. In the Sankey plot, each line represents a labeled neuron. **(B)** Heatmap (left) and Sankey plot (right) showing the distribution of the retrogradely labeled neurons in the anatomical subregions of IC and claustrum. **(C)** Representative spatial maps showing the spatial distribution of retrogradely labeled neurons (red) for each of the 14 projection targets in the anterior (top) and posterior (bottom) IC and adjacent regions. **(D)** Representative spatial maps showing the spatial distribution and cell identities of the retrogradely labeled neurons in the anterior (top) and posterior (bottom) IC and adjacent regions. The cells are color-coded by their transcriptomic identities.

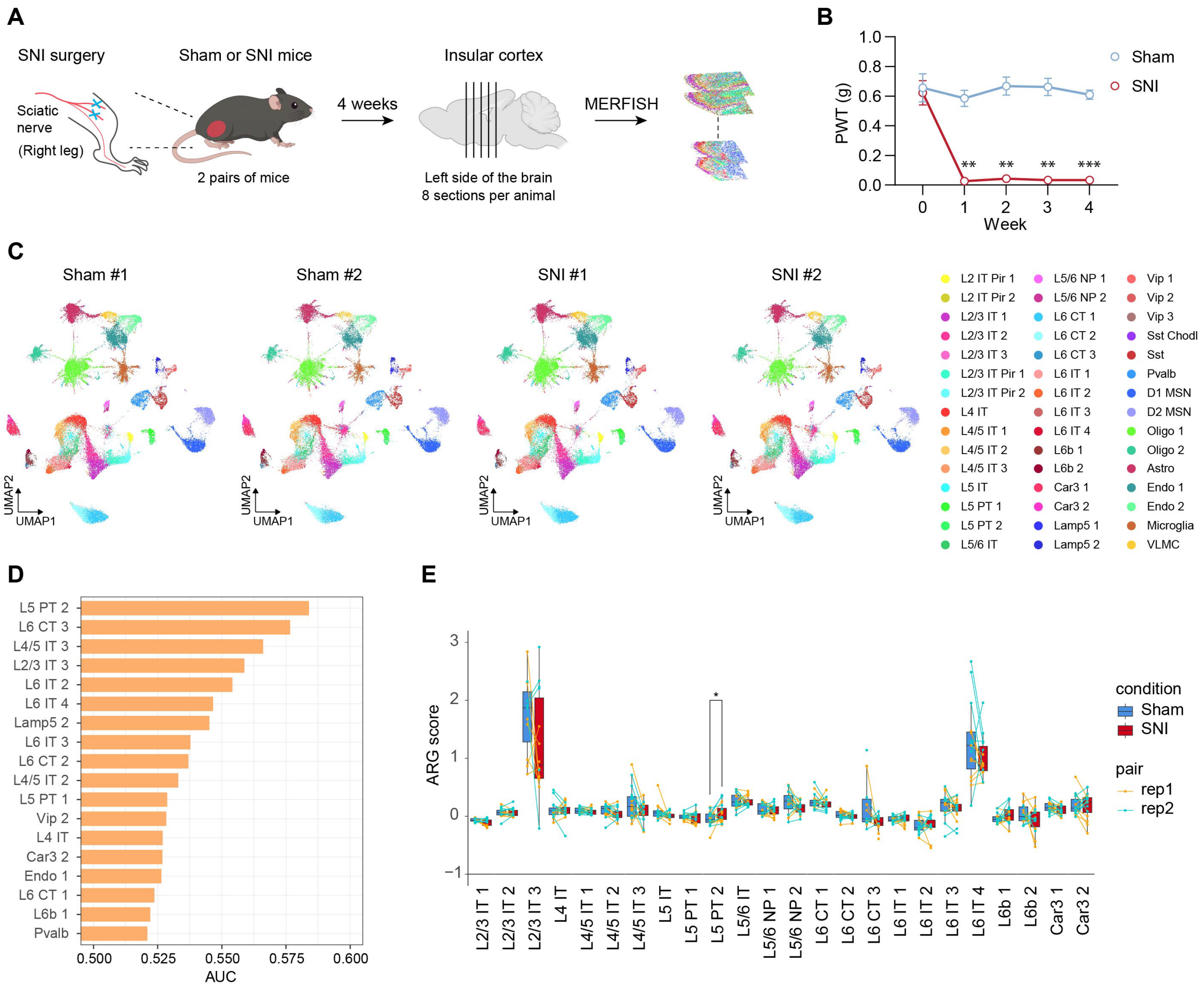

**Fig. S8. Transcriptomic impacts of chronic neuropathic pain on the IC. (A)** Diagram of the workflow to induce chronic neuropathic pain in mice and dissection of the IC for MERFISH analysis. **(B)** Mechanical allodynia of the mice subjected to SNI surgery compared to the control group revealed by von Frey test (n = 5 mice for each group). RM two-way ANOVA with Sidak's multiple comparison tests. **(C)** UMAP visualization of the cellular subtypes of the IC from mice with or without chronic neuropathic pain. The cells are color-coded by their identities. **(D)** Ranking of potential transcriptionally perturbed cell-types predicted by Augur. AUC shows the area under ROC curve of the predictions. **(E)** ARG scores of IC excitatory subtypes in control (blue) and pain (red) samples. Color of the paired dots indicates the paired animals.

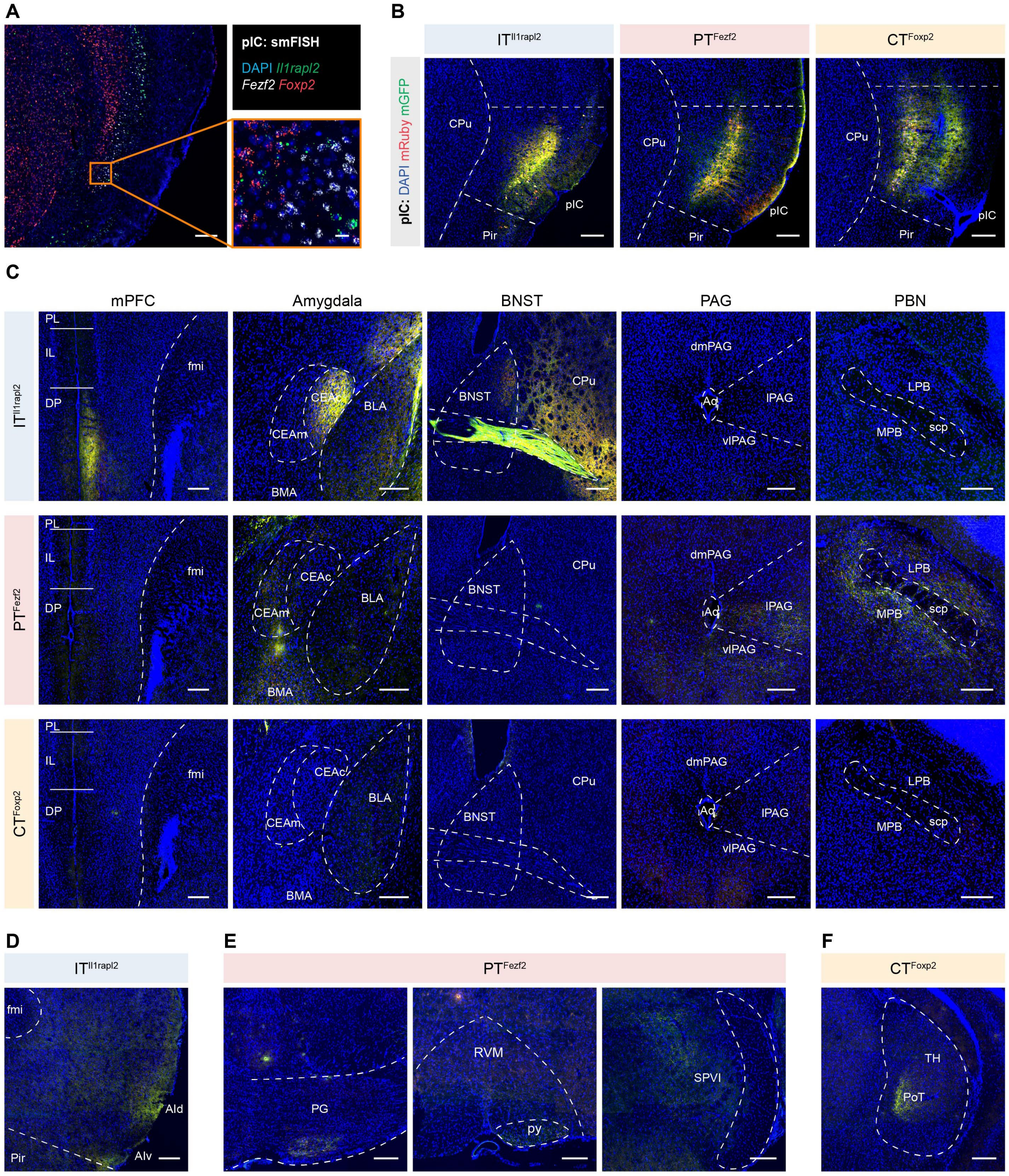

**Fig. S9. Anterograde tracing of the IT$^{Il1rapl2}$, PT$^{Fezf2}$ and CT$^{Foxp2}$ neurons in the pIC. (A)** Representative image of the smFISH for *Il1rapl2* (green), *Fezf2* (white) and *Foxp2* (red) in the pIC (left) and an enlarged image of the indicated region (right). Scale bars, 200 µm (left) and 20 µm (right). **(B)** Virus expression at the injection sites in pIC of the *Il1raple*-Cre, *Fezf2*-CreER and *Foxp2*-Cre mice. Scale bars, 200 µm. **(C)** Representative images showing the projections of the pIC IT$^{Il1rapl2}$ (top), PT$^{Fezf2}$ (middle), and CT$^{Foxp2}$ (bottom) neurons to the mPFC, Amygdala, BNST,

PAG and PBN in the three mouse lines. Scale bars, 200 μm. **(D)** Representative image showing the projections of the $IT^{Il1rapl2}$ neurons to the aIC. Scale bar, 200 μm. **(E)** Representative images showing the projections of the $PT^{Fezf2}$ neurons to the PG (left), RVM and pyramidal tract (middle) as well as SPVI (right). Scale bars, 200 μm. **(F)** Representative image showing the projections of the $CT^{Foxp2}$ neurons to the PoT. Scale bar, 200 μm. Abbreviations are in table S6.

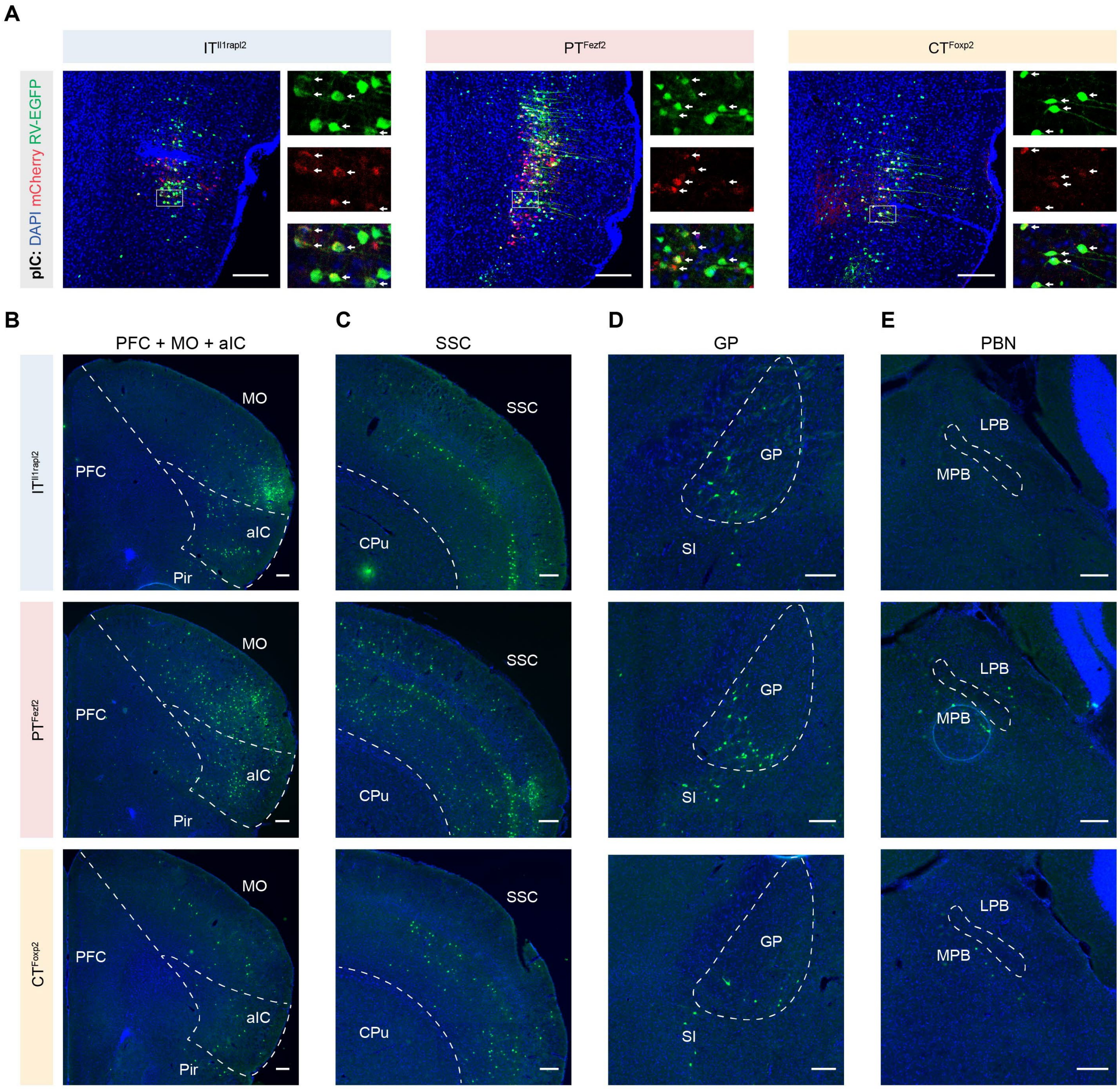

**Fig. S10. Monosynaptic retrograde tracing of the IT$^{Il1rapl2}$, PT$^{Fezf2}$ and CT$^{Foxp2}$ neurons in the pIC. (A)** Representative images showing the virus expression at the injection site in the pIC of the *Il1raple*-Cre (left), *Fezf2*-CreER (middle) and *Foxp2*-Cre (right) mice, and the enlarged images of the indicated regions showing the starter cells (arrows). Scale bars, 200 µm. **(B)** Representative images showing the retrogradely labeled monosynaptic input neurons (green) in the PFC, MO and aIC for the pIC IT$^{Il1rapl2}$ (top), PT$^{Fezf2}$ (middle), and CT$^{Foxp2}$ (bottom) neurons. Scale bar, 200 µm. **(C-E)** Same as **(B)** but for the SSC **(C)**, GP **(D)** and PBN **(E)**. Scale bars, 200 µm. Abbreviations are in table S6.

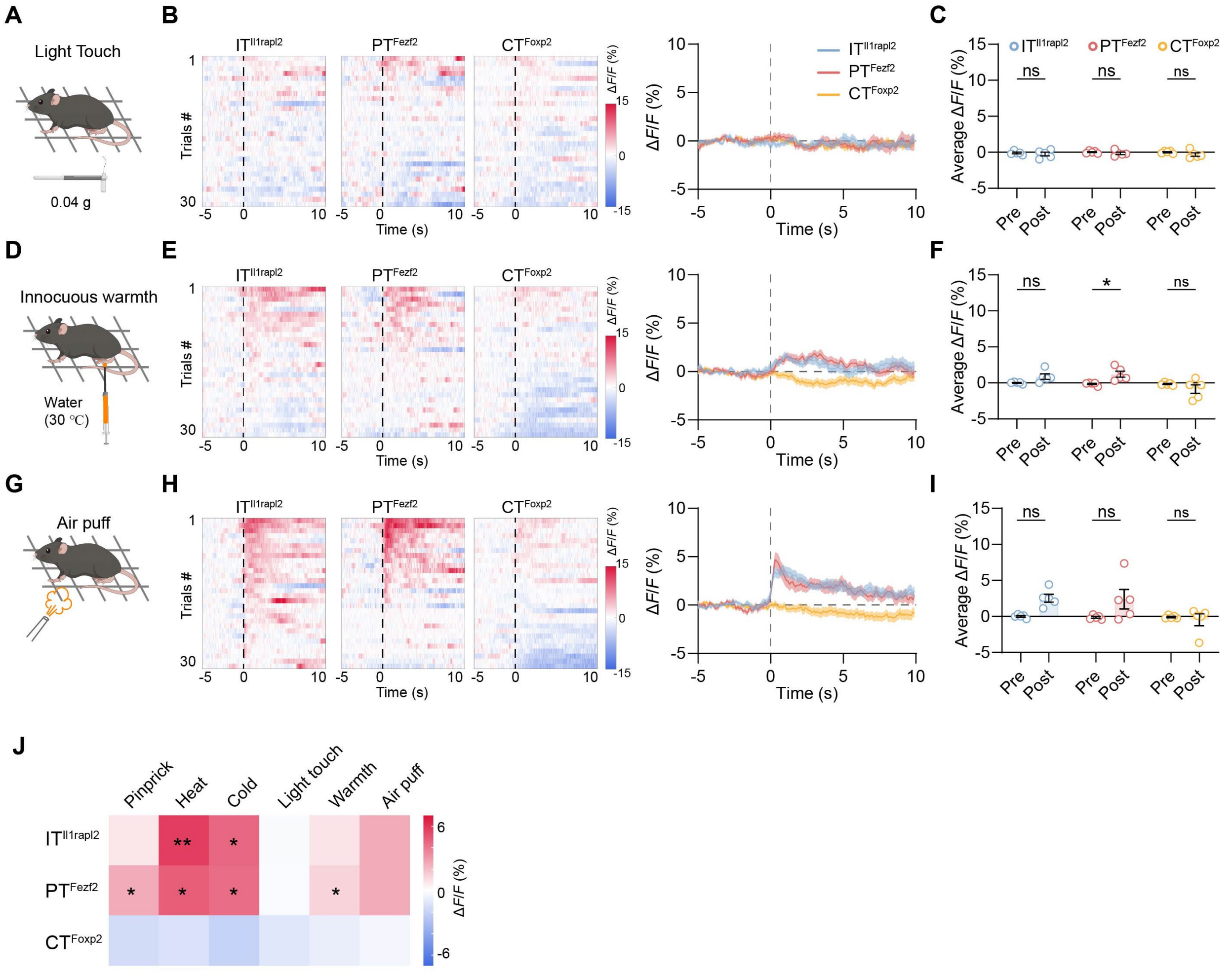

**Fig. S11. Neuronal activity of the $IT^{Il1rapl2}$, $PT^{Fezf2}$ and $CT^{Foxp2}$ neurons in the pIC in response to innocuous stimuli. (A)** Diagram of light touch stimulation to the mice. **(B)** Heatmaps (left) and averaged traces (right) showing the calcium activity of the $IT^{Il1rapl2}$ (blue), $PT^{Fezf2}$ (red) or $CT^{Foxp2}$ (yellow) neurons in the pIC during light touch stimulation. The dashed lines indicate the onset of light touch stimulation. The solid lines and shadow indicate the mean and s.e.m., respectively. n = 5 mice and 6 trials for each mouse. **(C)** Averaged ΔF/F (%) of the three neuron populations in the pIC 5 s before (pre) and after (post) the stimulation. **(D-F)** similar to **(A-C)** but for innocuous warmth stimulation with warm water (30 °C). **(G-I)** similar to **(A-C)** but for air puff stimulation. **(J)** Heatmap summary of the calcium activity of the three molecularly defined cell types in the pIC in response to noxious and innocuous stimulation. RM two-way ANOVA with Sidak's multiple comparison tests for **(C)**, **(F)** and **(I)**. *$P < 0.05$; ns, not significant.
